# Edge-based general linear models capture high-frequency fluctuations in attention

**DOI:** 10.1101/2023.07.06.547966

**Authors:** Henry M. Jones, Kwangsun Yoo, Marvin M. Chun, Monica D. Rosenberg

## Abstract

Although we must prioritize the processing of task-relevant information to navigate life, our ability to do so fluctuates across time. Previous work has identified fMRI functional connectivity (FC) networks that predict an individual’s ability to sustain attention and vary with attentional state from one minute to the next. However, traditional dynamic FC approaches typically lack the temporal precision to capture moment-by-moment network fluctuations. Recently, researchers have ‘unfurled’ traditional FC matrices in ‘edge cofluctuation time series’ which measure time point-by-time point cofluctuations between regions. Here we apply event-based and parametric fMRI analyses to edge time series to capture high-frequency fluctuations in networks related to attention. In two independent fMRI datasets in which participants performed a sustained attention task, we identified a reliable set of edges that rapidly deflects in response to rare task events. Another set of edges varies with continuous fluctuations in attention and overlaps with a previously defined set of edges associated with individual differences in sustained attention. Demonstrating that edge-based analyses are not simply redundant with traditional regions-of-interest based approaches, up to one-third of reliably deflected edges were not predicted from univariate activity patterns alone. These results reveal the large potential in combining traditional fMRI analyses with edge time series to identify rapid reconfigurations in networks across the brain.

## 1. Introduction

Our ability to selectively attend to task-relevant information varies across time. For example, while reading a book, you might periodically find yourself wondering if you have a message from a friend and end up checking your phone. You may have to jump pack a few paragraphs to find the place where you lost focus. Such attention fluctuations have been quantified with vigilance tasks, in which participants are tasked with detecting rare target stimuli during prolonged periods of watch (e.g., Mackworth 1948), and continuous performance tasks (CPTs), in which participants detect rare targets in a rapid stimulus stream (e.g., Robertson et al., 1997).

The gradual-onset continuous performance task (gradCPT; Esterman et al., 2013 Rosenberg et al., 2013; Fortenbaugh et al., 2015), a type of CPT in which image stimuli gradually fade from one to the next, was developed to assess sustained attention fluctuations while avoiding attention-capturing abrupt stimulus onsets (but see Jun & Lee, 2021). The gradCPT requires participants to make button presses in response to images from a frequently presented category (e.g., city scenes; 90%) while withholding responses to images from a rare target category (e.g., mountain scenes; 10%). Like other “not-X” CPTs (which require responses to frequent-category stimuli but not to rare targets), the gradCPT is a powerful tool for investigating sustained attention because it offers two methods for investigating attention fluctuations. First, participants’ responses to rare targets can be analyzed to assess successful withholding (reflecting engaged attention) or failures to inhibit the prepotent response (indicating an attention lapse). Second, the variability of response times to frequent-category images can be analyzed to produce a more high-frequency measure of attentional state known as the variance time course (VTC). Evidence suggests that more consistent response times (RTs) close to a person’s overall mean RT correspond to engaged attention. In contrast, more erratic responding (including both overly fast and slow responses) predict attention lapses and disengaged focus (Esterman et al., 2013; Rosenberg et al., 2013; Fortenbaugh et al., 2015; Kucyi et al., 2017; Jayakumar et al., 2023). Thus, fluctuating VTC values, calculated as the absolute deviation of each RT from the participant mean, are thought to reflect fluctuating sustained attention.

Analyses of the neural activity patterns underpinning sustained attention fluctuations have revealed seemingly counterintuitive results. When gradCPT RTs are consistent and performance is accurate, functional MRI activity in the default mode network (DMN)—typically associated with off-task and internally-directed thought and mind wandering (Buckner et al., 2009; 2019)—tends to rise above baseline (Esterman et al., 2013; Esterman et al., 2014; Kucyi et al., 2016; Kucyi et al., 2017; Fortenbaugh et al., 2018; Song et al., 2022). When RTs are erratic and performance suffers, fMRI activity in brain regions associated with top down control, such as regions of the dorsal attention network, tends to increase (Esterman et al., 2013; Esterman et al., 2014; Fortenbaugh et al., 2018; Kucyi et al., 2016; Kucyi et al., 2017; Song et al., 2022). This pattern of results may reflect practiced (Mason et al., 2007), less effortful performance during high attentional states and more effortful performance that relies on top-down control mechanisms during low attentional states (Esterman et al., 2013).

The activity approach has been informative but is limited in two ways. First, univariate bold activity has been shown to be a limited description of the signals present in the brain. For example, the feature-specific contents of working memory can be decoded from primary visual cortex during the delay period of a task via multivariate pattern analysis, despite no difference in the BOLD response between the delay period and the intertrial interval in those same regions (Serences et al., 2009). Second, the results are often examined at the network level, for example by examining activity changes in regions identified as part of the DMN, and are used to support claims about certain networks driving results. However, activation approaches cannot always speak to the functional configuration of networks. For example, activity in regions within a network may become more synchronized with increasing attention, without changing in overall activation.

An alternative approach has involved studying functional networks more directly by relating individual functional connectivity (FC) matrices, computed using time series data collected while participants performed attention tasks or rest in the scanner, to attention function (e.g., Kessler et al., 2016; Poole et al., 2016; Wu et al., 2020; Kucyi et al., 2021; Yoo et al., 2022). For example, one approach identified sets of functional connections, or edges, whose strength predicts individual performance (*d’*) on the gradCPT (Rosenberg et al., 2016). A connectome-based predictive model based on strength in these edges can generalize across task and rest scans, and even predict attention-related clinical symptoms, suggesting that it captures a key neural signature of sustained attentional differences. In addition, FC matrices from small time windows can predict ongoing sustained attention performance across a scale of tens of seconds to minutes (Kardan et al., 2022; Rosenberg et al., 2020). However, one’s attentional state can change rapidly over seconds (e.g., during a brief lapse), and windowed FC approaches are limited by the window size used, preventing them from capturing these high-frequency fluctuations in attention. This contrasts with the relative temporal granularity of GLM fitting to univariate activity: GLM approaches are limited by the time course of the hemodynamic response function, which peaks around 5s and is resolved after around 20 seconds.

In summary, two separate fMRI approaches have related univariate activity and functional connectivity to sustained attention. The former has examined the activation of the DMN and control networks during high and low attentional states. The latter has examined the relationship between the strength of edges within the functional connectome and attentional performance across individuals and across time. However, neither of these methods directly tests for the type of rapid network reconfigurations that may underlie fluctuations in attention. Traditional activation-based approaches are limited in the sorts of changes they can identify across conditions or individuals, while functional connectivity approaches are limited in their temporal granularity and are underpowered for capturing high-frequency changes. These limitations aren’t restricted to research on sustained attention, but apply to any cognitive phenomenon that can be understood at the network level and which varies at a high frequency.

Recent advances in network neuroscience provide the possibility of combining the FC perspective with the ability to capture high-frequency fluctuations via GLM fitting. Specifically, researchers have begun to “unfurl” the functional connectivity matrix into its time point-by-time point contributions, called edge cofluctuations (Faskowitz et al., 2020, Zamani Esfahlani, 2020). These edge cofluctuations reflect estimates of covariance between the pair of brain regions comprising the edge at each moment in the time series. The mean of these estimates across time is Pearson’s *r* between the two regions’ time series, the most common FC metric. Edge cofluctuations capture higher-frequency changes in covariance between brain regions than traditional dynamic FC approaches, which incorporate information across multiple observations in a time window.

Here, we combine the analytical methods for univariate activity with a FC perspective to ask how rapid network reconfigurations relate to high-frequency fluctuations in sustained attention. To address this question, we examined two independent gradCPT fMRI datasets. We first examine the replicability of standard univariate activity approaches (Fortenbaugh et al., 2018). Following this, we fit standard GLMs to edge time series to capture attentional lapses, and correlate the VTC with edges. In almost all cases, we identified a set of edges significantly deflected in both datasets. A large proportion of these edges are not predicted from just univariate activity. Furthermore, edges that reliably fluctuate with attentional performance significantly overlap with an existing FC model of sustained attention (Rosenberg et al., 2016). Finally, we compare significance thresholding approaches with varying assumptions and consider how the edge response functions may differ from the canonical hemodynamic response function. We propose that fitting GLMs to edge time series offers a powerful lens into rapid network reconfigurations underlying cognitive operations.

## 2. Methods

### 2.1. Dataset 1

To assess the reproducibility of edge-behavior associations, we examined fMRI data from two independent datasets collected while participants performed the gradual-onset continuous performance task (gradCPT; Esterman et al., 2013), which measures sustained attention and inhibitory control. The first dataset, dataset 1, was described in Rosenberg et al. (2016).

#### 2.1.1. Participants

Dataset 1 included 25 participants with normal or corrected-to-normal vision (13 female, 12 male, age 18-32 years, mean age 22.7 years). One participant was excluded due to a task lag in one run, leaving 24 participants for analysis. Participants gave written informed consent in accordance with the Yale University Human Subjects Committee and were paid for their participation.

#### 2.1.2. Experimental design

Participants completed three fMRI runs of the gradCPT. Runs consisted of four three-minute blocks (225 800-ms trials/block). Blocks were separated with 32-second rest blocks. Participants were instructed to respond to city scenes (90% of trials) and to withhold responses to mountain scenes (10%) with an emphasis on accuracy. An iterative algorithm assigned key presses to trials as described previously (Esterman et al., 2013). The algorithm first assigned unambiguous button presses, which occurred after image *n* was 80% cohered but before image *n + 1* was 40% cohered. Ambiguous presses were then assigned to an adjacent trial if one of the two had no response, to the closest trial if both had no response, or to an adjacent city trial if the other adjacent trial was a mountain. If multiple presses could be assigned to a trial, the fastest response was selected. On rare occasions, the original algorithm incorrectly assigned responses from the first trial of a block to the last trial of the preceding block, resulting in long response times. We recalculated RTs from the original dataset.

#### 2.1.3. fMRI acquisition parameters

MRI data were collected at the Yale Magnetic Resonance Research Center on a 3T Siemens Trio TIM system using a 32-channel head coil. Before task runs, an anatomical magnetization prepared rapid gradient echo (MPRAGE) was collected. Task runs consisted of 824 whole-brain volumes acquired using a multiband echo-planar imaging (EPI) sequence: repetition time (TR) = 1000 ms, echo time (TE) = 30 ms, flip angle = 62°, acquisition matrix = 84 x 84, in-plane resolution = 2.5 mm^2^, 51 axial-oblique slices parallel to the ac-pc line, slice thickness = 2.5 mm, multiband = 3, acceleration factor = 2. A 2D T1-weighted image with the same slice prescription as the EPI images was collected for registration.

#### 2.1.4. fMRI data preprocessing and parcellation

Data were preprocessed using BioImage Suite (Joshi et al., 2011) and custom Matlab scripts (Mathworks) as described previously (Rosenberg et al., 2016). The first 8 TRs of each run were excluded from analysis. Motion was corrected using SPM8 (http://www.fil.ion.ucl.ac.uk/spm/software/spm8/). Linear and quadratic drift, mean signal from cerebrospinal fluid, white matter, and gray matter were regressed from the data along with a 24-parameter motion model (6 motion parameters, 6 temporal derivatives, and their squares). Data were then temporally smoothed with a zero mean unit variance Gaussian filter. Functional MRI data were then parcellated using the 268-node Shen atlas (Shen et al., 2013).

### 2.2. Dataset 2

Dataset 2 was described in Yoo et al. (2022). Although rest and other task runs were also collected, we only analyze gradCPT runs here. Dataset 2 is available at https://nda.nih.gov/edit_collection.html?id=2402.

#### 2.2.1. Participants

127 participants performed the gradCPT during two MRI sessions separated by approximately two weeks on average. The original authors excluded 35 participants for excessive head motion (>3 mm maximum head displacement or >.15 mm mean framewise displacement), fewer than 120 TRs after censoring, low data quality via visual quality assurance, or task performance 2.5 *s.d.* from the group mean in both sessions, leaving 92 participants (60 females, 32 males, ages 18-35 years, mean age 22.79 years). Exclusion criteria were applied to each run, and participants were excluded if they had no run for any of the five run types (three tasks, one rest, and one move-watching). Of these, 65 participants had behavioral performance measures in both gradCPT sessions. We analyze data from the 58 participants in this group with fewer than 10% of censored whole-brain volumes. Participants gave written informed consent in accordance with the Yale University Human Subjects Committee and were paid for their participation.

#### 2.2.2. Experimental design

Participants completed one 10-minute gradCPT run containing 740 trials in a single block in each MRI session. The task parameters are the same as those used in dataset 1.

#### 2.2.3. fMRI acquisition parameters

FMRI data were collected at the Yale Magnetic Resonance Research Center and the Brain Imaging Center at Yale on 3T Siemens Prisma systems using a 64-channel head coil. Before task runs, an anatomical MPRAGE scan was collected. Task runs consisted of 600 whole-brain volumes acquired using a multiband EPI sequence: TR = 1000 ms, TE = 30 ms, flip angle = 62°, acquisition matrix = 84 x 84, in-plane resolution = 2.5 mm^2^, 52 axial-oblique slices parallel to the ac-pc line, slice thickness = 2.5 mm, multiband = 4, acceleration factor = 1.

#### 2.2.4. fMRI data preprocessing and parcellation

Data were preprocessed using Analysis of Functional NeuroImages (AFNI version 17.2.07) as described previously (Yoo et al., 2022). The first 3 TRs of each run were excluded from analysis. Volumes were censored if they contained outliers in more than 10% of voxels or if the Euclidean distance of the head motion parameter derivatives were greater than 0.2 mm. Data were then despiked, slice-time corrected, and motion corrected. Mean signal from cerebrospinal fluid, white matter, and gray matter were regressed from the data along with 6 motion parameters, 6 temporal derivatives, and their squares. Data were then aligned to the MPRAGE anatomical image and normalized to MNI space. FMRI data were then parcellated using the 268-node Shen atlas (Shen et al., 2013). Thirteen nodes were excluded from analysis in at least one participant due to imperfect acquisition of the voxels within the region, predominantly within the frontoparietal and motor networks (5 nodes each), and within the medial-frontal (2 nodes) and subcortical-cerebellar (1 node) networks.

### 2.3. Analyses

Analyses were conducted in Python 3.10.2 using a combination of nilearn (https://nilearn.github.io/stable/index.html), scikit-learn (Pedregosa et al., 2011), and custom scripts. Our main analyses were preregistered (https://osf.io/ftzr4/) and the code is available (github.com/henrymj/edge_GLM). Post-hoc analyses and deviations from the preregistration are noted as such below. Throughout this paper, we use the term first-level model to refer to within-individual analyses, second-level model to refer to assessments of significance at the group level within each dataset, and third-level to refer to assessments of overlap between datasets. Visualization of connectivity matrices were built using seaborn’s heatmap plotting functions (Waskom, 2022)

#### 2.3.1. First-level analyses: ROI-based activation

We first asked how univariate fMRI activity in the 268 Shen atlas ROIs covaried with attention fluctuations following analyses described in Fortenbaugh et al. (2018). Fortenbaugh et al. (2018) examined how changes in activity reflected fluctuations in attention and attention lapses during the gradCPT, replicating original results from Esterman et al. (2013) in a larger sample (*n* = 140; one 8-min gradCPT run per participant). They examined attentional lapses by contrasting pre-trial and trial-evoked activity to erroneous responses to rare mountain scenes (commission errors, CEs) against correct withholding of responses (correct omissions, or COs). To characterize attention fluctuations, they correlated fluctuations in response time with activation time series. These analyses contextualize edge-based results by indicating the reproducibility of each analysis and demonstrating what edge-based analyses add beyond traditional ROI-based approaches^1^.

##### 2.3.1.1. Event-based contrasts

We built a first-level model with the following regressors:

1. Incorrect presses to rare mountain scenes (commission errors, CEs), which occur when participants fail to inhibit their prepotent response and indicate a lapse in attention;
2. correct withholding of responses to rare mountain scenes (correct omissions, COs), indicating that participants successfully inhibited their prepotent response and thus are likely in a focused, high-attention state;
3. incorrect failure to respond to common city scenes (omission errors, OEs), which may result from a variety of sources.

As in Fortenbaugh et al., (2018), we did not model correct responses to common city scenes (correction commissions) due to their high frequency.

Stick regressors for each modeled trial type were convolved with the SPM hemodynamic response function (HRF) model and its temporal derivative to create the design matrix. Because nuisance regressors (motion, drift, white matter, global signal) were regressed out during preprocessing, they were not included in these models. To better equate the baseline period between dataset 1, which included rest breaks between task blocks, and dataset 2, which did not contain breaks, we included a single block break regressor per run which lasted the duration of each break in dataset 1.

We fit each participant’s time series for each run to the design matrix using nilearn’s autoregressive (AR) model, which whitens the data according to the autoregressive covariance structure in the time series. Censored timepoints in dataset 2 were interpolated from adjacent timepoints, and an additional impulse regressor was included in the model for each censored timepoint. We also confirmed that excluding censored timepoints and fitting an ordinary least squares model did not qualitatively change results. Statistics were averaged across runs within each subject.

Using this model, we computed the four main contrasts of Fortenbaugh et al. (2018): CE against baseline, CO against baseline, OE against baseline, and CO – CE. One run from one participant in dataset 1 is excluded, as they did not have any correct omissions for that run; the participant’s remaining two runs were kept. One participant in dataset 2 is excluded from these contrasts as they did not make any omission errors, leaving a sample size of 57 in dataset 2 for these analyses.

##### 2.3.1.2. Correlating activity with RT variability

The contrasts above ask how activity differs between trial types. To characterize how activity fluctuates with a continuous measure of attentional state, we separately correlated activation time series with each run’s variance time course (VTC). The VTC is computed using the response times (RTs) on correct frequent-category trials (i.e., correct presses to city trials or correct commissions) as described previously (Rosenberg et al., 2013; Esterman et al., 2013). RTs were *z*-scored and converted to absolute values. Trials missing responses (OEs and COs) were linearly interpolated from the values of the two surrounding trials. This time course was then smoothed using a Gaussian kernel with a full-width at half-maximum (FWHM) of 9 trials, producing the VTC. The VTC was computed separately within each of the four blocks per run in dataset 1. In dataset 2, each run consisted of a single block. To account for the lag of the HRF, we shifted the VTC by 6 seconds following Fortenbaugh et al. (2018).

We then correlated the VTC with residuals of each ROI’s BOLD-signal time series after fitting the event-based GLM described above. Time points from the between-block breaks were excised from the residuals in dataset 1. Correlation values were converted to *z*-values using Fisher’s transformation to be assessed using our second- and third-level paradigms below.

##### 2.3.1.3. Identifying precursors of attention lapses

The above analyses asked how ROI activity corresponds to attention lapses, task events, and continuous fluctuations in sustained attention. We next asked whether activity preceding rare mountain trials *predict* upcoming correct omissions vs. commission errors. To do so we replicated the trial-precursor analysis of Fortenbaugh et al. (2018), applying the details of their whole-brain analysis. ROI time courses were linearly interpolated in time to the onset of each trial. For each CO or CE, the preceding timepoints from –4.8s (6 trials) to the onset of the trial were averaged. These preceding averages were averaged across trials of the same type, and a difference map (CO minus CE) was generated for each participant. As in the contrast analyses above, one run from one participant in dataset 1 was excluded, as they did not produce any correct omissions for that run; the participant’s remaining 2 runs were kept.

#### 2.3.2. First-level analyses: Edge cofluctuations

We next asked how edges rather than activity in a single region alone covary with attention. To this end, we computed edge cofluctuation time series for a given pair of nodes (ROIs), *i* and *j*, following (Faskowitz et al., 2020). Specifically, each ROI’s time series vector (*i* or *j*) was z-scored, and the elementwise product between the two standardized vectors was computed. Mathematically, the edge time series *e_ij_* is computed as:

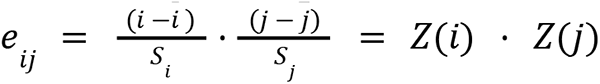

Where x̄ is the mean and *S_x_* is the standard deviation across the time series for node *x*:

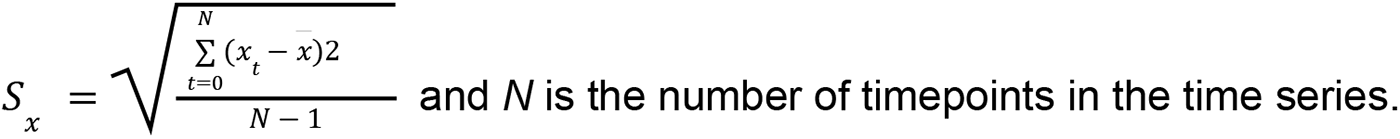

We replicated all activation analyses with edge (i.e., cofluctuation) time series as described in detail below.

##### 2.3.2.1. Relating cofluctuations to attention lapses and salient events

We first asked how edges covary with attention lapses and fluctuations. To do so, we built first-level models with regressors for the following trial types: commission errors (CEs), correct omissions (COs), and omission errors (OEs). As in our ROI activity analyses, we computed the four main contrasts of Fortenbaugh et al. (2018): CE against baseline, CO against baseline, OE against baseline^2^, and CO – CE. In addition to these four contrasts, we included a CO + CE contrast, which, given the structure of the task, is equivalent to a no-go over baseline contrast, or an oddball (i.e., rare mountain trial) vs. baseline contrast. This analysis allows us to identify whether certain task-based events (attention lapses, correct inhibition, no-go, and oddball processing) are accompanied by high-frequency changes in the functional interactions between brain regions.

##### 2.3.2.2. Relating cofluctuations to RT variability

We next explored whether edges varied systematically with the VTC, the continuous measure of attention computed from RT deviance (Esterman et al., 2013; Rosenberg et al., 2013). As in our ROI-activity analyses, we correlated the lag-shifted VTC with edge time series after excising timepoints from the between-block breaks in dataset 1, and converted the correlations to *z*-values using Fisher’s transformation for group-level assessment.

In addition to this time-shifted smoothed VTC analysis, we asked whether a more traditional parametric regressor analysis (Esterman et al., 2013) might be better powered than the original correlational approach. Rather than smoothing the VTC with a gaussian kernel, we convolved the non-smoothed VTC with the SPM HRF function and its temporal derivative and included these in a model with trial event regressors, similar to that described above. We included an event regressor for each of the four trial types (correct commissions, correct omissions, commission errors, and omission errors). A block break regressor was included for dataset 1. We fit this model to the edge time series and computed the contrast VTC over baseline^3^.

##### 2.3.2.3. Identifying precursors of attention lapses in edge cofluctuations

As in our ROI activity analyses, we asked whether edge cofluctuations preceding rare mountain trials predict upcoming correct omissions vs. commission errors. First, edge time courses were linearly interpolated in time to the onset of each trial and then averaged across the preceding timepoints from -4.8s to the onset of each rare mountain trial. We produced a difference map (CO – CE) of these average preceding values for each participant.

#### 2.3.3. Second-level significance testing via permutation

We tested the significance of each first-level analyses at the group level using two different family-wise error rate corrections: (1) max-T for both ROI and edge analyses, and (2) the network-based statistic (NBS; Zalesky et al., 2010) for edge analyses. These methods rely on different assumptions to assess statistical significance. We provide both results so that other researchers interested in applying edge-based GLMs can use the one best suited for their research question.

Max-T controls for type-1 error by producing a null distribution of the maximum test statistics (or minimum *p*-values) across random permutations of the data, to which the observed edge test statistics can be compared. It makes no assumptions about the relationships between tests (in other words, about the relationship between edges, or ROIs), so it can be applied to both ROI activity and edge cofluctuation first-level maps.

NBS is a graph-specific procedure that also controls for type-1 error by constructing a null distribution across random permutations of the data, but it makes the additional assumption that edges that are significantly different from the null will be interconnected in subcomponents of the full graph. This is analogous to the assumption that significant voxels will be spatially autocorrelated which underlies cluster correction in traditional fMRI contrasts. Thus, the null distribution for NBS is formed of maximum subcomponent sizes across permutations. In both cases, we used the sign-flipping paradigm from the randomise algorithm (Winkler et al., 2014) to randomly permute edges or ROIs.

First, we fit a second-level model to the first-level outputs and obtained an observed test statistic for each edge. Because this work is exploratory and we have no directional hypotheses, we chose to use the F-statistic. We then ran 10,000 permutations of the second-level model by randomly flipping the signs of the intercept column (i.e., multiplying each participant’s vector of edge statistics by 1 or -1) and computing new F-statistics for each edge.

To build the max-T distribution, we recorded the largest F-value. To build the NBS distribution, we thresholded the permutation’s second-level graph to edge-behavior associations with *p* < .01 and recorded the size of the largest fully connected component, defined by the number of edges in the component.^4^ For edges, the max-T and NBS null distributions were computed in parallel. The observed second-level maps were then thresholded using these two null distributions, producing two sets of results. For max-T, observed ROI-behavior or edge-behavior associations were thresholded to those with F-statistics greater than the 95^th^ percentile of the null distribution. For NBS, observed edge-behavior associations were thresholded to those with *p* < .01, and any components larger than the 95^th^ percentile of the NBS null distribution were retained.

#### 2.3.4. Third-level analyses for edges: between dataset comparisons via permutation

The reproducibility of the ROI results can be assessed via comparison with previous literature, such as Esterman et al. (2013) and Fortenbaugh et al. (2018). In contrast, there is no existing work to compare the edge results against. Therefore, we rely on comparisons between our two independent datasets to assess the reliability and reproducibility of these results. To assess whether the edges identified as significant were consistent across datasets, we relied on a permutation test with slight differences in implementation to account for the different assumptions of our two second-level paradigms.

To assess overlap in the max-T results, we recorded the number of overlapping significant edges between our two datasets. Across 10,000 permutations, we shuffled the locations of significant edges in both datasets and recorded the number of overlapping edges.

To assess overlap in the NBS results, we repeated the above process, but we shuffled node locations rather than edges, as NBS makes assumptions about the connections within components. Shuffling nodes therefore randomizes edge locations while preserving the structure of the observed components.

For both max-T and NBS, a result was considered to have significant consistency if the number of observed overlapping edges was larger than the 95^th^ percentile of the null distribution of overlapping edges.^5^

### 2.4. Assessing dataset-level overlap with edges underlying individual differences in attentional control

This work aims to identify edges associated with sustained attentional performance by looking for differences from the null (i.e., no association) at the group level. In contrast, previous work has identified edges associated with attentional performance by examining individual differences in performance. Rosenberg et al. (2016) applied connectome-based predictive modeling (CPM; Finn et al., 2015; Shen et al., 2017) to predict individuals’ gradCPT *d’*. CPM uses a cross-validation approach to identify edges whose strengths are most positively or negatively correlated with attentional performance. The set of most positively correlated edges are termed the high-attention network (HAN), and the set of most negatively correlated edges are termed the low-attention network (LAN). CPM identifies the high- and low-attention networks in a training set of data, then fits a simple linear model to that training set to predict *d’* for a given individual as the weighted combination of their average high-attention edge strength and their average low-attention edge strength. This model is then tested on held out data.

In this exploratory analysis, we asked whether the edges identified by our group-level approach overlapped with those identified via CPM. To do this, we assessed the overlap between preregistered high- and low-attention network edges and the second-level results from three analyses isolating attentional control: 1) the CO-CE contrast, 2) the CO-CE precursor analysis, and 3) correlations with the VTC. We focused on overlap with NBS-based second-levels, as the max-T results produced very sparse graphs. Because the CPM edges were identified in dataset 1, we examined the results for each dataset separately and focus on significant overlap between the CPM edges and dataset 2 results.

We assessed significant overlap between second-levels and CPM edges in nearly the same manner that we assessed significant overlap between datasets (see Methods section 2.3.4). We first recorded the observed overlap with the following assumptions. We assumed that positively deflected edges for CO-CE (for either the contrast or precursor analysis) should map onto HAN edges and negatively deflected edges should map onto LAN edges. In contrast, we expect HAN and LAN edges to map onto the edges correlated with the VTC differently.

Increasingly positive VTC values reflect increasingly erratic responding, associated with reduced attention; edges which are positively correlated with the VTC reflect increased synchrony with reducing attention. Thus, we expect that edges which are positively correlated with the VTC should map onto LAN edges, and edges which are negatively correlated with the VTC should map onto HAN edges. Then, across 10,000 permutations, the second-level maps were permuted at the node level to preserve the subcomponent structure, and overlap with the CPM edges were recorded. Observed overlap was assessed against this null distribution.

### 2.5. Comparing observed edge results to ROI-based predictions

Do edge-based GLMs reveal new information about brain-behavior relationships, or are they redundant with the results of traditional ROI-based GLMs? To ask this question, we investigated whether the results based on edge cofluctuation time series contained information distinct from that contained in ROI time series and traditional analyses. In each dataset, we compared our second-level edge-based results against theoretical edge second-levels predicted from ROI activity. To predict the edge-based results from the ROI results, we converted each first-level ROI map to a “theoretical edge map” using the following formula:

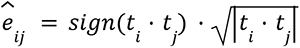

where *t* is the *t*-score for a given node (*i* or *j* in this example).

This formula captures a few straightforward predictions. First, if both nodes are deflected (i.e., have non-zero *t*-scores), their edges are likely to also be deflected. Second, if both nodes are deflected in the same direction, this will likely lead to a positive deflection in the edge, and if the nodes are deflected in opposite directions, this will likely lead to a negative deflection in the edge.

We analyzed these maps using the second-level tests described above and compared contrasts with significant overlapping edges between datasets in both the ROI-based and

edge-based maps. We predicted that there may be pairs of regions whose activation time series are not significantly deflected from baseline in response to task events, yet whose edge cofluctuation time series were.

### 2.6. Estimating hemodynamic response function of edge time series

Though the timing of edge fluctuations should still be constrained by the speed of the hemodynamic response function, it is possible that the standard SPM HRF provides a poor description of how edge time series are deflected by events. In this post-hoc analysis, to estimate the hemodynamic response function of edges in response to events, we employed finite impulse response modeling and *k*-means clustering. We performed this estimation procedure on each participant’s first session in dataset 1, so that the estimated response functions may be applied to the second session as well as all of dataset 2, should these response functions differ in form from the HRF.

We fit our trial-type model to both the ROI and edge time series for each participant’s first run of dataset 1. Rather than convolving the trial-type stick regressors with the HRF, we used 31 separate impulse regressors per trial-type (one for each TR of the first 30 seconds after an event). To increase power, we focused on parameter estimates for the CO+CE contrast, which includes the most data.

We then ran a group-level test for each ROI or edge at each timepoint, and averaged across participants. Including only those ROIs or edges with a value significantly different from 0 for at least one time point produced similar HRFs to using all ROIs or edges. To avoid separating response functions with the same structure but different directions, each time series was multiplied by the sign of its largest magnitude value (i.e., if an ROI or edge’s largest value was negative, the whole time series was flipped).

We then clustered the ROI or edge time series using *k*-means clustering. We found an optimal number of clusters by testing all values of K between 2 and 30 (inclusive) and identifying the elbow in the distortion function across Ks (yellowbrick;

https://www.scikit-yb.org/en/latest/api/cluster/elbow.html). We took the mean, or centroid, of each cluster to be a representative HRF.

## 3. Results

### 3.1. Activation patterns replicate previous work on sustained attention

In both datasets, the three baseline contrasts produced a similar pattern of results to that reported in Fortenbaugh et al. (2018) (**Figure 1**; compare with their Figure 4). Correct omissions were associated with greater activity in task-positive frontal and parietal regions previously associated with vigilant attention (e.g., Langner & Eickhoff, 2013) and lesser activity in regions of the default mode network often associated with task-irrelevant or internally directed thought (Buckner et al., 2009; 2019). As in Fortenbaugh et al. (2018), lateral visual cortex activity increased with correct omissions whereas medial visual cortex activity decreased. Commission errors were associated with similar activity patterns, potentially reflecting attention capture by and processing of rare-target images. In addition, both trial types likely involve attempts at inhibitory control processes. Omission errors, which occurred rarely, were again accompanied by similar activity patterns including increased activity in the middle frontal gyrus, inferior parietal lobule, and insula and decreased activity in ventromedial prefrontal cortex. To facilitate visual comparison with the results of Fortenbaugh et al. (2018), we projected the ROI activation results onto the fsAverage surface mesh from FreeSurfer, using nilearn’s functions *vol_to_surf* and *plot_surf_stat_map*. Vertices were assigned the value of the nearest voxel. To get a sense of the ROI boundaries on this surface projection, we mapped the ROI labels to a set of 6 circularly repeating, highly contrasting colors, and surface projected this mapping in the same manner as well (**Figure S1**).

**Figure 1.**
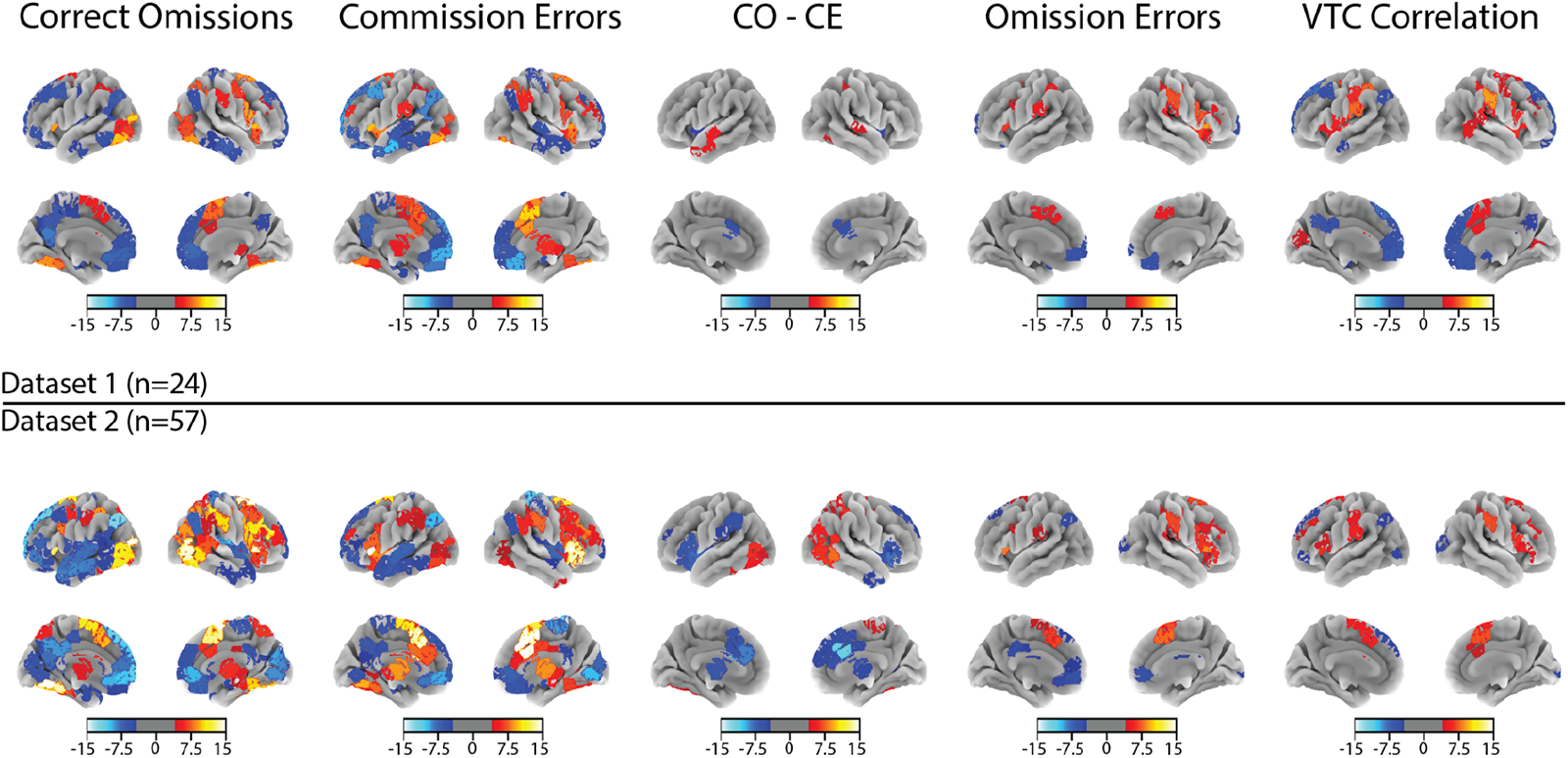
ROI-based activation contrasts for trial-type events. Each column reflects one of the four main event-based contrasts of Fortenbaugh et al. (2018), or the correlation between the variance time course (VTC) and ROI time series. Each row reflects the dataset-specific second-levels, which were thresholded using a max-T permutation testing approach (see Methods section 2.3.3). Note that because these analyses were conducted at the ROI level and then surface projected, there are shared contours to the activation sites. For a reference of the ROI boundaries when surface projected, see **Figure S1**.

Contrasting correct omissions with commission errors (CO-CE) in dataset 2 replicated Fortenbaugh et al. (2018). Successful inhibition was associated with more activity in subcortical and cerebellar regions and visual and inferior parietal areas presumably driven by greater attention-to-task. Unsuccessful inhibition, on the other hand, was associated with greater medial prefrontal and dorsal anterior cingulate cortex and insula activity, potentially reflecting error processing (Dali et al., 2023). A sparser CO-CE map in dataset 1 suggests that this smaller sample may be underpowered to detect effects consistent in Fortenbaugh et al. (2018) and dataset 2, although this interpretation of the null effect is speculative.

Associations between continuous attention fluctuations and ROI activity replicated the overall pattern of results observed in Fortenbaugh et al. (2018) (**Figure 1**; compare with their Figure 7). In both datasets, the smoothed, lag-shifted VTC was negatively correlated with activity in regions of the default mode networks, replicating previous associations between higher DMN activity and in-the-zone, more practiced sustained attention task performance (Mason et al., 2007; Esterman et al., 2013; Esterman et al., 2014; Kucyi et al., 2020; Song et al., 2022). This relationship is hypothesized to reflect the fact that, during “in-the-zone” attentional states, good performance may be less effortful and need not rely on the suppression of DMN activity (Esterman et al., 2013). In contrast, higher activity in task-positive frontal and parietal regions was positively correlated with RT variability, potentially reflecting more effortful top-down attention during erratic periods of performance (Esterman et al., 2013; Kucyi et al., 2017; Fortenbaugh et al., 2018).

Pre-trial activity did not significantly vary before correct omissions vs. commission errors in dataset 1. In dataset 2, greater frontal and parietal activity preceded correct omissions whereas greater ventromedial prefrontal cortex activity preceded commission errors (**Figure S4**; compare with Fortenbaugh et al. [2018] Figure 5). This aligns with previous work suggesting that, while “task-positive” frontoparietal activity may decrease and “task-negative” DMN activity may increase during in-the-zone attentional performance, attention lapses can occur when frontoparietal activity is too low or DMN activity is too high (Esterman et al., 2013).^6^ Notably, however, both datasets tested here have fewer participants than Fortenbaugh et al. (2018) and may be underpowered for detecting attention-lapse precursors.

### 3.2. Edge-GLM reveals significant high-frequency fluctuations in connectivity across networks

Associations between ROI activity and sustained attention performance generally replicated previous associations between in-the-zone performance and increased DMN and decreased frontoparietal activity while providing tentative support for the idea that *too much* DMN activity or *too little* frontoparietal activity can cause lapses. Errors caused by these lapses are accompanied by greater activity in regions associated with error processing, and rare targets in general are associated with more task-positive and less DMN activity.

Together these findings and the body of work they replicate motivate the prediction that changes in interactions within and between *networks* of regions—not just individual regions themselves—vary with attentional performance. This hypothesis is not directly testable with traditional dynamic functional connectivity analyses that rely on measuring region interactions in windows of BOLD-signal time series. Thus, we applied GLMs to edge time series to assess relationships between attention task events and performance at the level of region pairs.

To do so, we applied the same analyses described previously to edge rather than BOLD-signal time series. We asked whether edge time series reliably covaried with task or cognitive events and assessed 5 contrasts reflecting attention lapses/inhibitory control failures (CE vs. baseline), successful inhibition (CO vs. baseline and CO-CE), failures to respond (OE vs. baseline), and rare-target processing (CO+CE vs. baseline) using traditional trial-type regressors (**Table 1** and **Figures 2** and **S2-3)**.

**Table 1.**
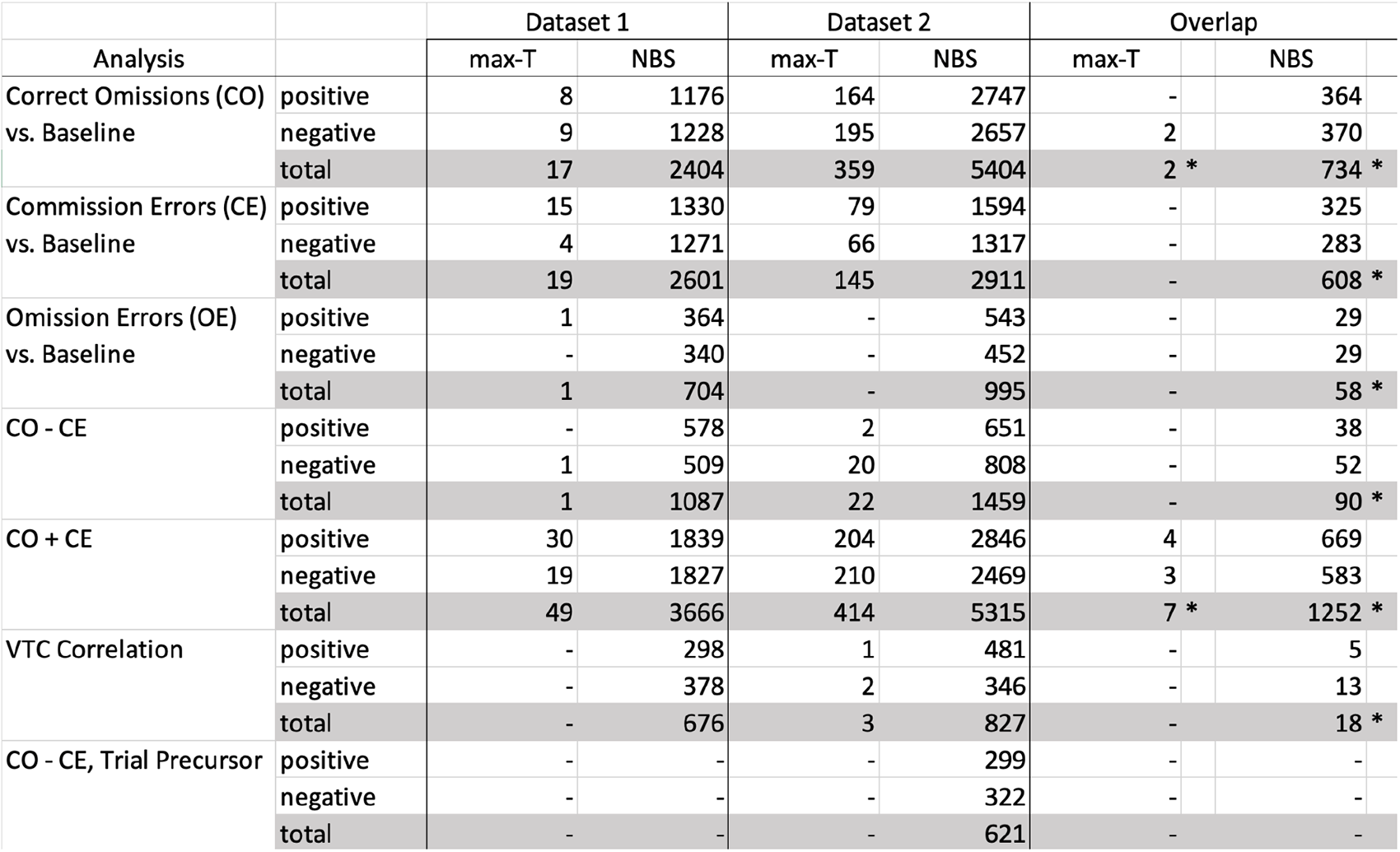
Quantitative summary of the significant edges for each analysis. For each dataset, the number of positive and negative edges are listed for each second-level method. The far right column reflects the number of overlapping edges between datasets for each analysis and second-level method. Asterisks (*) reflect a significant amount of overlap between the datasets.

| Analysis |  | Dataset 1 |  | Dataset 2 |  | Overlap |  |
| --- | --- | --- | --- | --- | --- | --- | --- |
|  |  | max-T | NBS | max-T | NBS | max-T | NBS |
| Correct Omissions (CO)<br>vs. Baseline | positive | 8 | 1176 | 164 | 2747 | - | 364 |
|  | negative | 9 | 1228 | 195 | 2657 | 2 | 370 |
|  | total | 17 | 2404 | 359 | 5404 | 2 * | 734 * |
| Commission Errors (CE)<br>vs. Baseline | positive | 15 | 1330 | 79 | 1594 | - | 325 |
|  | negative | 4 | 1271 | 66 | 1317 | - | 283 |
|  | total | 19 | 2601 | 145 | 2911 | - | 608 * |
| Omission Errors (OE)<br>vs. Baseline | positive | 1 | 364 | - | 543 | - | 29 |
|  | negative | - | 340 | - | 452 | - | 29 |
|  | total | 1 | 704 | - | 995 | - | 58 * |
| CO - CE | positive | - | 578 | 2 | 651 | - | 38 |
|  | negative | 1 | 509 | 20 | 808 | - | 52 |
|  | total | 1 | 1087 | 22 | 1459 | - | 90 * |
| CO + CE | positive | 30 | 1839 | 204 | 2846 | 4 | 669 |
|  | negative | 19 | 1827 | 210 | 2469 | 3 | 583 |
|  | total | 49 | 3666 | 414 | 5315 | 7 * | 1252 * |
| VTC Correlation | positive | - | 298 | 1 | 481 | - | 5 |
|  | negative | - | 378 | 2 | 346 | - | 13 |
|  | total | - | 676 | 3 | 827 | - | 18 * |
| CO - CE, Trial Precursor | positive | - | - | - | 299 | - | - |
|  | negative | - | - | - | 322 | - | - |
|  | total | - | - | - | 621 | - | - |

We first assessed edges that consistently deflected with correct omissions. Activation results revealed that COs were associated with more frontoparietal and lateral visual cortical activity. The edge-based GLM revealed edges associated with these networks and others.

Max-T permutation testing identified two reliable edges (where “reliable” refers to significant deflection in both datasets; probability of overlap given the number significant edges in each dataset is *p*=.004; **Figure S3**), both of which were negative deflections between the motor and visual association networks. NBS permutation testing identified 734 edges common to both datasets (*p*=1/10,001). These consisted of negative deflections between the motor and visual, and subcortical-cerebellar networks but positive deflections between motor and medial frontal, frontoparietal, and default mode networks. Regions in these higher-order association networks also showed positive deflections with visual networks in response to COs. Negative deflections within the medial frontal, frontoparietal, default mode, and motor networks—and a similar number of positive and negative deflections within the subcortical-cerebellar network—also accompanied COs. Thus, COs tended to be accompanied by increased cofluctuation between higher-order association networks and motor and visual regions and decreased cofluctuation between motor, visual, and subcortical-cerebellar networks. This may reflect attention capture by rare target stimuli and/or the engagement of top-down attentional control leading to successful inhibition.

Unsuccessful inhibition (CE against baseline) identified 0 overlapping edges using the max-T approach and 608 overlapping edges (*p*=1/10,001) using the NBS approach. As in the activation based results, the overall pattern was similar to that of COs, with some differences. For example, more edges within the medial frontal and frontal parietal networks were negatively deflected whereas more edges within the subcortical-cerebellar network were positively deflected. The medial frontal network also showed more negative deflections with the frontoparietal and default mode networks and positive deflections with the motor and visual networks.

We isolated successful inhibitory control and sustained attention by comparing correct omissions against commission errors (CO-CE). Activation analyses found increased activity in subcortical, cerebellar, visual, and inferior parietal regions in response to successful inhibition, but increased activation in control and error processing related regions in response to unsuccessful inhibition. Max-T permutation testing on edges did not identify overlap between the datasets. NBS permutation testing identified 90 overlapping edges (*p*=1/10,001), which primarily involved regions of the subcortical-cerebellar and medial frontal networks. Successful inhibition was associated with more positive deflections within medial frontal and visual association networks and more negative deflections within the subcortical-cerebellar network. Successful inhibition was also associated with more negative deflections between the medial frontal network and the frontoparietal, subcortical-cerebellar, motor, and visual association networks and more positive deflections between the subcortical cerebellar network and every other network except for the medial frontal network. Speculatively, this pattern of results may suggest that greater medial frontal network segregation (more positive within- and negative between-network cofluctuation) and greater subcortical-cerebellar network integration (more negative within- and positive between-network cofluctuation) accompanies successful relative to unsuccessful inhibition.

We contrasted omission errors (incorrectly withholding responses to common city trials) against baseline. Activation patterns found increased activity in the middle frontal gyrus, inferior parietal lobule, and insula, and decreased activity in the ventromedial prefrontal cortex. While Max-T based permutation testing did not identify any overlapping edges, NBS based permutation testing identified 58 overlapping edges (*p*=1/10,001). The edges consisted mostly of positive deflections within the motor network and negative deflections distributed across the brain networks. Most of the negatively deflected edges were associated with the medial frontal network, both within-network and with the default mode network and subcortical cerebellar network, or were associated with connections between the motor network and the subcortical cerebellar and the visual II network.

Lastly, we examined rare trials against baseline (CO+CE; **Figure S3**). Max-T identified 7 overlapping edges (p=1/10,001). These consisted of negative deflections between the motor network and both the visual association and subcortical-cerebellar networks, along with positive deflections between the subcortical-cerebellar network and both high-level vision networks and the frontoparietal network. NBS permutation testing identified 1252 overlapping edges (p=1/10,001), with patterns of deflections roughly combining those seen separately in the CO against baseline and CE against baseline contrasts.

**Figure 2.**
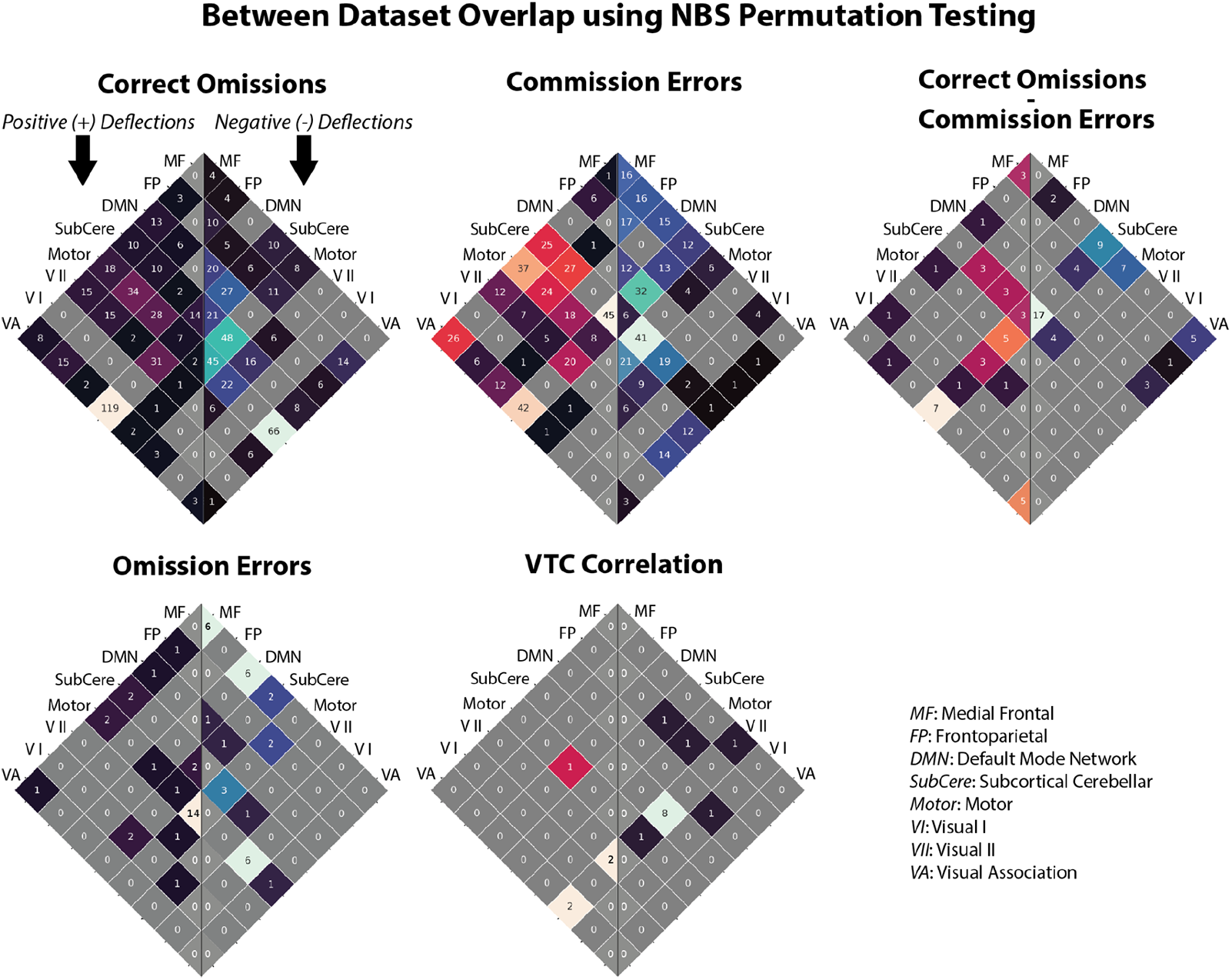
Cross-network distribution of reliable (i.e., significant in both datasets) edges from trial-type contrasts, based on second-levels computing using NBS permutation testing. Each of the first four matrices reflects the overlap of significant edges for a given contrast: Correct Omissions (CO), Commission Errors (CE), rare probe trials (CO+CE), and successful inhibitory control (CO-CE). The final matrix reflects edges that were positively or negatively correlated with the variance time course (VTC). Each heat map shows the number of edges in a given set of network-network connections that were significant for a given analysis. The left side of each heatmap shows the number of edges with positive deflections, while the right side of each heatmap shows the number of edges with negative deflections.

### 3.3. Edges covary with continuous fluctuations in attentional states across time

We investigated whether edge fluctuations track continuous fluctuations in attentional states in two ways. For each, we began by computing the VTC, a measure of attentional focus (Esterman et al., 2013; Fortenbaugh et al., 2018). In the first analysis, we replicated Fortenbaugh et al. (2018)’s analysis by correlating each edge time series with the VTC, shifted by 6 seconds to account for hemodynamic lag. We transformed the correlations into *z*-scores via Fisher’s transformation and applied our second-level and between-group analyses. While max-T permutation testing did not identify an overlapping set of edges, NBS-based permutation testing identified a significant overlapping set in both the correlation approach (*n*=18, *p*=.002). Most correlations with the VTC were with edges within and between the visual II network. Edges within this network and with the visual association network were positively correlated with VTC values, while edges with the subcortical-cerebellar network were negatively correlated. There was some evidence of the involvement of higher-order association regions: a DMN-motor edge was positively correlated with the VTC while two frontoparietal-motor and frontoparietal-subcortical cerebellar edges were negatively correlated. This suggests that, as attention waned, cofluctuations between frontoparietal, vision, and motor production networks became increasingly negative, while cofluctuations within vision networks and with the DMN became increasingly positive. In contrast, increases in attention are associated with increased functional connectivity between frontoparietal, vision, and motor production networks. The second planned parametric regressor analysis did not identify any significant edges using max-T or any significantly large components using NBS in either dataset.

### 3.4. Edges may not vary preceding successful versus unsuccessful responses

We examined edge time series preceding correct omissions and commissions errors in response to rare mountain trials (Fortenbaugh et al., 2018). We averaged the preceding 4.6s before each CO and CE trial, and took the difference between trials types for each participant. Max-T based permutation testing did not identify significant differences in either dataset. NBS based permutation testing also did not identify any significantly different subcomponent in dataset 1, but did identify a subcomponent containing 621 edges in dataset 2. These included both positive and negative deflections within the subcortical-cerebellar and motor networks as well as between these networks and higher-order association and visual networks (**Table 1**; **Figure S4**).

### 3.5. Edges correlated with attentional fluctuations support individual performance predictions

We assessed whether edges that were significantly deflected in three key analyses isolating attentional control overlapped with edges previously identified as underlying individual differences in sustained attention and attentional control using connectome-based predictive modeling (CPM; Rosenberg et al., 2016). We selected the NBS-thresholded second-levels of 1) the CO-CE contrast, 2) the CO-CE precursor analysis, and 3) correlations with the VTC. Because the CPM edges were originally identified in dataset 1, we focus on overlap with second-level results in dataset 2.

Neither of the CO-CE analyses (contrast and precursor analyses) identified a significantly overlapping set of edges (contrast: n=32, *p*=.28; precursor: n=16, *p*=.17). In contrast, there was significant overlap between the edges that correlated with the VTC and the attention networks identified via CPM (n=55, *p*=1/10,001; **Figure 3**). This overlap analysis was built on the assumption the high-attention network (HAN) edges should map onto edges negatively correlated with the VTC and low-attention network (LAN) edges should map onto edges positively correlated with the VTC (see Methods section 2.4). Only edges matching these mappings were considered during the permutation test. Of the 58 edges identified as overlapping between the VTC second level and the CPM edges, 55 matched the mapping. A chi-squared test confirmed that the two discretizations (HAN/LAN and positive/negative correlations) were dependent (χ^2^_Yates_ = 42.68, *p*=6*10^-11^), supporting our assumption.

HAN/negatively correlated edges were largely distributed across edges between frontal networks and motor/visual networks (**Figure 3**). In contrast, LAN/positively correlated edges were distributed across subcortical cerebellar, motor, and visual association networks.

**Figure 3.**
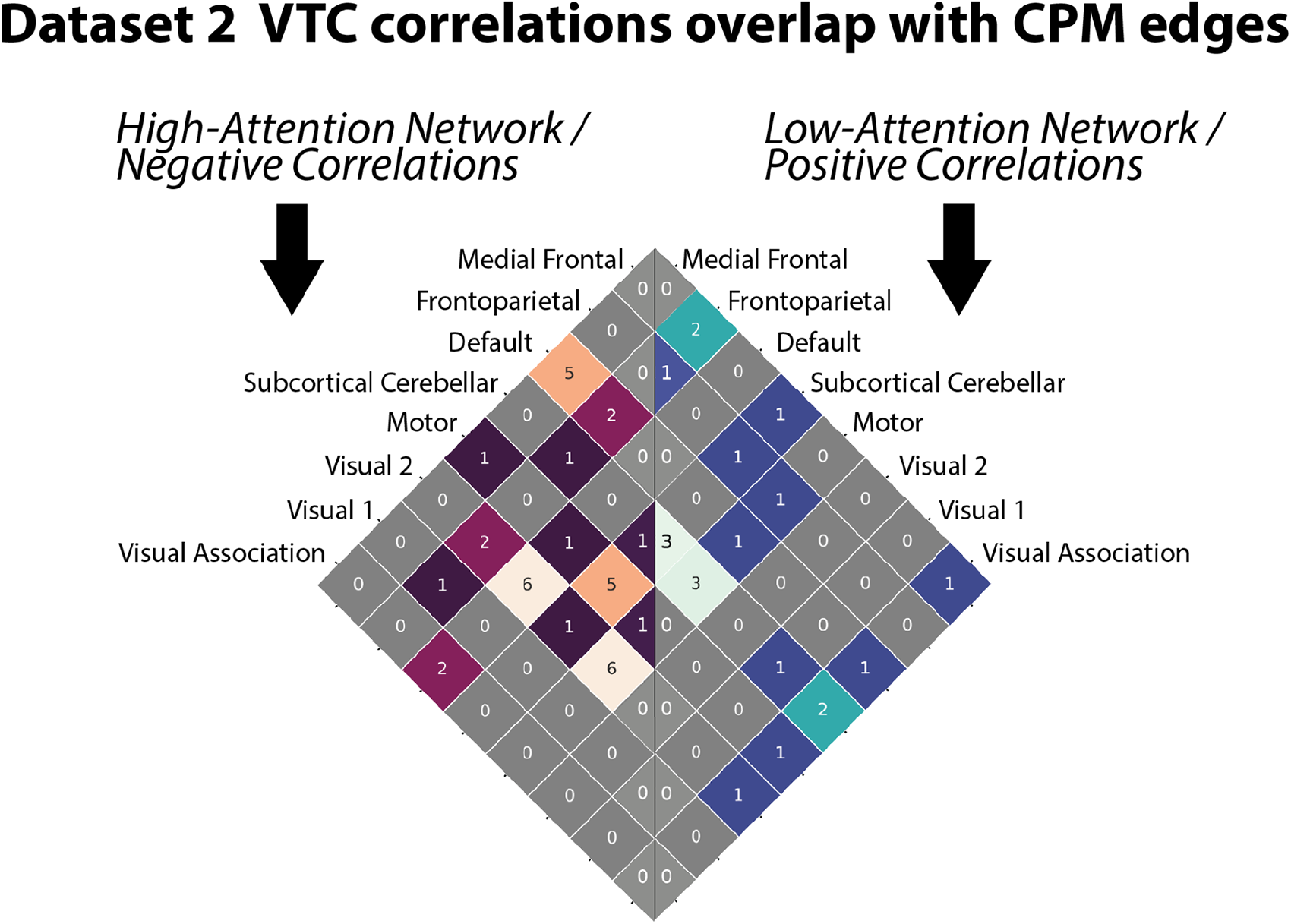
Overlap of edges which significantly correlated with the VTC in dataset 2, and which contributed to predictions of individual differences in performance in dataset 1. Each cell of the matrix represents the number of significant edges between two networks (or within a network, if lying along the diagonal). The left half reflects the count of high-attention network edges that overlapped with edges that were significantly anti-correlated with the VTC. The right half reflects the count of low-attention network edges that were significantly positively correlated with the VTC.

### 3.6. Edge fluctuations can be systematically mispredicted by ROI-based predictions

To test whether the information captured by our edge time series analysis was redundant with information from traditional ROI activation analyses, we fit the ROI-time-series with the contrast models introduced above. We then converted the first-level results to predicted edge results by multiplying the *t*-scores for each pair of ROIs and taking the signed square-root (see Methods section 2.5). We applied our second- and third-level comparisons to these predicted edge values and identified edges *predicted* to be non-significant in either dataset but in reality significant in both datasets.

Across the five event-based contrasts within our NBS-based second-levels, 19-33% of all edges that were significant in both datasets were not predicted to be significant in either dataset based on their activation alone. This is not because the ROI-based predictions produced more sparse maps than the observed edge maps. In each dataset and in each of the five contrasts, as many or more edges were identified as significant when predicted from ROI activation maps compared to being actually observed. With the Max-T permutation paradigm, this increase ranged from 0% (an equal number of edges were identified as significant) to 4,317% (17 significant edges observed vs 751 predicted) in dataset 1, and from 772% (22 vs 192) to 1,361% (359 vs 5,248) in dataset 2. The increase was smaller when using the NBS permutation paradigm, but still substantial (around 100-200%). Thus, edge-based GLMs are related to but not redundant with activation analyses.

### 3.7. Edge responses are similar to traditional hemodynamic response functions

The above results indicate that we are able to capture changes in edge time series using models assuming the canonical HRF. However, it is possible that not all edge responses are described by the HRF, meaning that model designs may not be optimized. To test this, we estimated ROI and edge responses to rare trials versus baseline using finite impulse repulse (FIR) modeling (see Methods section 2.6).

Nine clusters were identified for succinctly capturing the ROI HRFs produced by our contrast (**Figure 4**). Six of the centroids, accounting for 165 ROIs (61.6% of all ROIs), were shaped as the canonical HRF, and varied mostly in amplitude and timing (peaking between four and six seconds). The remaining three centroids, accounting for 103 ROIs (38.4%) were shaped similarly to the canonical HRF, but decayed slowly after peaking, rather than dropping roughly in proportion to the activation rate. Two of these three centroids, accounting for 84 ROIs, had a somewhat late peak (8 and 10s). ROI response functions similar to these centroids would likely be partially matched by the canonical HRF, especially when modeled with its temporal derivative as an additional regressor, so we may assume that the canonical HRF is roughly appropriate for all ROIs.

**Figure 4.**
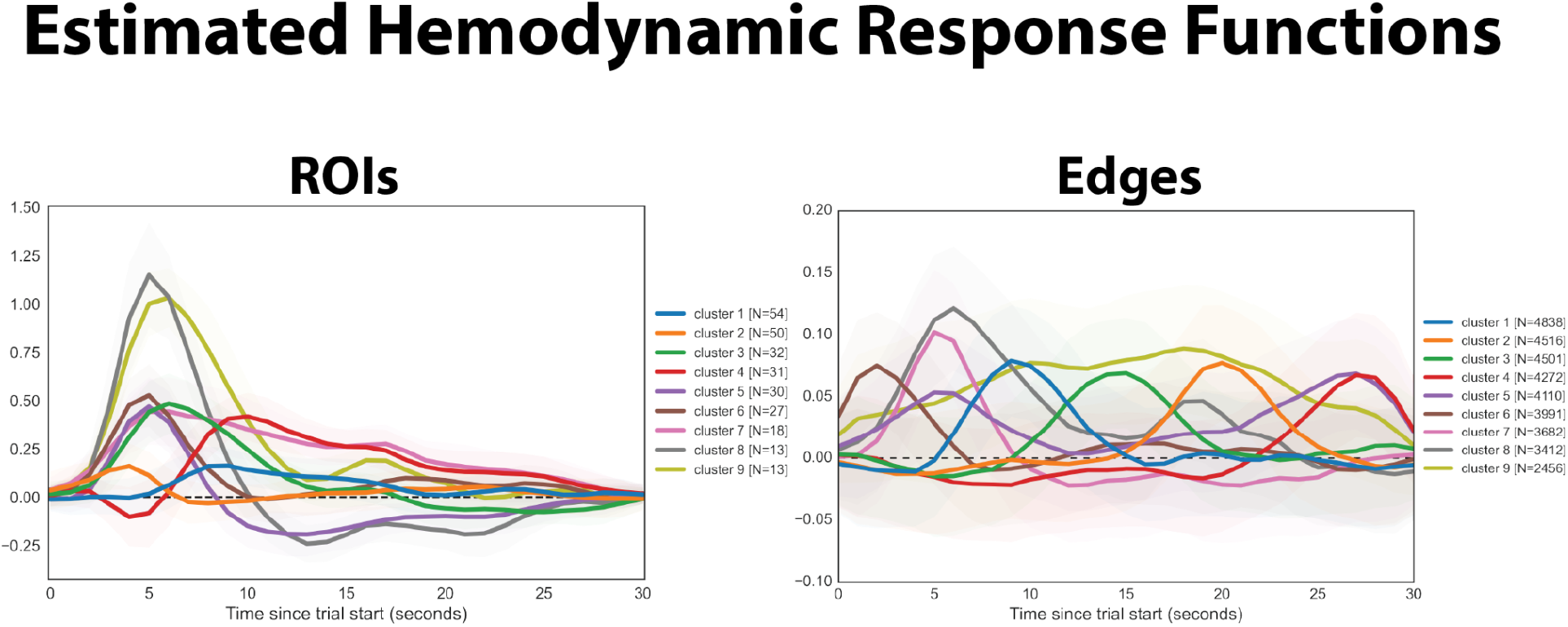
Estimated hemodynamic response functions from univariate ROI activity and edge time series. The responses for the 30 seconds following rare probe trials were estimated for each ROI and edge using finite impulse response modeling. Fits were performed for the first session of each subject in dataset 1, and averaged across subjects before being clustered using k-means. The centroid (mean) of 9 clusters are presented for each time series type, ordered by cluster size. Shaded regions reflect the bootstrapped 95% confidence intervals of the means.

Nine clusters were also identified for succinctly capturing the edge-HRFs produced by our contrast (**Figure 4**). Four of these, accounting for 15,923 edges (44.5% of all edges) appeared to match the canonical HRF with slight shifts in amplitude and timing (peaking between 2–9 seconds). Three of the remaining five centroids, accounting for 13,289 edges (37.1%) were also shaped similar to the canonical HRF, but with larger lags in their timing (peaking between 14–27 seconds). The final two centroids were not of the canonical HRF shape. One centroid, accounting for 4,110 edges (11.5%), contained two distinct peaks, one at five seconds, and one at 27 seconds. Edge response functions similar to this centroid would likely be partially matched by the canonical HRF, which peaks around 5 seconds and returns to baseline around 20 seconds after stimulus onset. The final centroid, accounting for 2,456 edges (6.9%), slowly rose, peaking around 18 seconds, then slowly fell. Thus, the canonical HRF, along with its temporal derivative, likely provided a strong-to-partial match for 44.5-56% of edges.

It is important to note that, though the canonical HRF is only appropriate for a smaller subset of edges compared to ROIs, this should only produce false negatives among edges with non-canonical response functions, not false positives. This suggests that the number of edges that were not predicted from univariate activity above may be an underestimate.

## 4. DISCUSSION

In this work, we examined whether traditional fMRI GLM analyses can be applied to edge cofluctuation time series to capture high-frequency fluctuations in attention. We assessed the replicability of results across two independent datasets in which participants performed the gradCPT, which enables examination of attention both at the event level and via continuous measures across time. In general, activation results replicated previous empirical findings (Esterman et al., 2013, Fortenbaugh et al., 2018). Cofluctuation results extended the activation results by identifying new sets of edges that covary with attention and overlap with networks that predict individual differences in sustained attention.

To assess the robustness of previous activation-based results and contextualize our edge-based results, we replicated the univariate activity analyses of Fortenbaugh et al. (2018). Contrasts against baseline all produced results aligning with Fortenbaugh et al. (2018). Correctly withheld responses to rare probe trials (correct omissions or COs), commission errors (CEs), and both together produced similar patterns of activity. These trials were associated with increased activity in regions associated with vigilant attention and decreased activity in the default mode network. They were also associated with increased lateral visual cortex activity and decreased medial visual cortex activity. This aligns well with a meta-analysis of 11 Go/No-go fMRI studies contrasting No-go and Go trials which implicated the bilateral putamen, middle occipital gyrus, insula, inferior parietal lobule, pre-SMA, middle frontal gyrus, the right superior and inferior frontal gyri, and the left fusiform gyrus (Simmonds, Pekar & Mostofksy, 2008). Erroneous withholding on frequent non-probe trials (omission errors or OEs) was also associated with regions connected to error processing including the inferior parietal lobule, insula, middle frontal gyrus, and ventromedial prefrontal cortex.

Replicating results of Fortenbaugh et al. (2018), successful withholding of responses (CO – CE) was associated with greater activity in subcortical, cerebellar, visual, and inferior parietal areas and lesser activity in the medial prefrontal cortex, dorsal anterior cingulate cortex, and insula activity in dataset 2 (but not dataset 1; **Figure 1**). To our knowledge, few studies have contrasted successful and unsuccessful inhibition directly, but one study which examined these conditions separately found increased cuneus, caudate and inferior frontal activity for successful stopping, and increasing anterior cingulate and insula activity with unsuccessful stopping (Chevrier, Noseworth & Schachar, 2007), both of which have previously been associated with error monitoring and adjustments of control (Ramautar, et al., 2006, Shenhav, Cohen, & Botvinick, 2016, Dali et al., 2023). Our CO – CE contrast then likely captures regions contributing to appropriate motor execution or withholding (subcortical, cerebellar, and inferior parietal areas) during successful inhibition, and error processing and control adjustments (medial PFC, dACC, insula) following unsuccessful inhibition.

Results also replicated associations between continuous fluctuations of attention (measured by the variance time course, or VTC) and ROI activity, revealing negative correlations between the VTC and default mode network activity and positive correlations between the VTC and task-positive frontal and parietal region activity. Results did not fully replicate pretrial differences between COs and CEs. There were no significant pretrial activity differences in dataset 1, though in dataset 2 the patterns of activity aligned with previous work suggesting that lapses occur when frontoparietal activity is too low or DMN activity is too high (Esterman et al., 2013). Overall, these activation contrasts replicated previous work and identified clusters of regions implicated in subprocesses relevant to the gradCPT, including visual processing, motor execution, error processing, control adjustments and top down attentional modulation.

The only two univariate analyses that did not qualitatively replicate prior results in both datasets were the CO-CE and pre-CO vs. pre-CE contrasts. It is perhaps not surprising that the CO-CE contrast and trial precursor analysis produced sparser and non-significant results, respectively, in dataset 1. Participants correctly omit responses to rare (10%) mountain trials roughly 70-80% of the time and make commission errors the remaining 20-30%. This means that the CO-CE contrast relies on a difference between measures based on 7-8% and 2-3% of trials. In addition, the trial precursor results have smaller effect sizes than the standard contrast approach (compare the *t*-values of Figure 4d and 5b in Fortenbaugh et al. [2018]). Therefore, it’s likely that these differences require relatively large samples to be reliably identified. Dataset 1 consisted of only 24 participants, which contrasts with the 58 participants of dataset 2 and 140 participants of Fortenbaugh et al. (2018). The component identified by NBS permutation testing in dataset 2 was also the smallest component identified in any analysis, which could also reflect a lack of power, although it is also possible that the component involved is just smaller than those involved in the other analyses. Larger sample sizes should allow us to better capture both smaller components and components with smaller effect sizes in the future.

While the univariate results can be interpreted in terms of networks, they cannot capture changes in network connectivity that occur without a change in overall activation and are thus limited. To address this limitation, we applied the above fMRI analyses to edge time series to ask whether changes in the cofluctuation of pairs of regions or networks are involved in changes in sustained attention. These edge-based results both aligned with and diverged from the univariate activation results (**Figure 2**). Contrasts with sparse univariate results also produced sparse edge results. In addition, though not captured by this overlap analysis, dataset 1 always produced sparser edge results than dataset 2 (**Figure S2**), as it did for the univariate results.

These reliable edges were distributed across the brain at the network level, though this distribution was not uniform. Many of the edges connected networks identified by the activation contrasts, specifically the subcortical-cerebellar, frontal, and default mode networks. The edges within and between these networks were variably deflected, both matching with predictions (e.g., positive deflections within the medial frontal network for correct omissions vs commission errors), and extending beyond the previous literature (e.g., that edges connecting the medial frontal with motor networks are mostly negatively deflected in the same contrast). How networks interact with one another has only been minimally predicted by the previous literature, which instead focuses on the relative activation of regions within a subset of networks as a proxy for the network’s engagement during the task. Deflections between frontal and subcortical-cerebellar regions and regions in the motor and vision networks emphasize the integration of processing from sensory input all the way to response execution during successfully sustained attention, and point to the need to examine network connectivity more directly.

Edges that correlated with VTC produced striking results. First, reliable edges were only found in networks associated with sensory processing and motor-production, namely the visual, motor, and subcortical-cerebellar networks (**Figure 2**), rather than association networks such as the frontal networks and the DMN. In addition, those significantly correlated edges in dataset 2 also significantly overlapped with edges identified using connectome-based predictive modeling (CPM) in dataset 1 to identify individual differences in gradCPT performance (**Figure 3**). A few features of this result are worth noting. First, the edge approach identifies edges that consistently vary with the VTC across individuals. In contrast, the CPM approach relies on variance across the group. Although this did not need to be the case, the same edges that track attention fluctuations at the group level also scale with individual differences in overall performance. Second, though the edges which were reliable across datasets were identified in sensory and motor-production networks, the edges overlapping between dataset 2 and the CPM edges also included many connections with the medial frontal and frontoparietal networks. This suggests again that dataset 1 may have been underpowered and failed to capture deflections in these frontal networks, previously implicated in sustained attention (e.g., Fortenbaugh et al., 2017).

Of note, we found that 19-33% of the reliable edges identified via contrast analyses were not predicted from univariate ROI activity, despite ROI activity predicting deflections in more edges for each contrast than were actually identified. Therefore, these edge analyses are extracting reliable and *novel* information that can provide a view into rapid network reconfigurations underlying fluctuations in cognitive processes.

Though this work identified reliable and novel edges while making minimal assumptions about how edge cofluctuations may differ from univariate activity, there are many options for optimizing this workflow and increasing the power of these analyses. First, we found that the HRF is appropriate for at least a subset of edges. Other work has also used a canonical HRF function to examine edge time series “events”, high amplitude moments that explain a large amount of variance in the connectomes between individuals, and found moderate correlations between a convolved movie-event time series and edge events time series (Tanner et al., 2022). It is worth noting that the edge events time series is a derivative of the edge time series, computed from the root sum square of all edges. The correlation between the convolved movie-event time series and the edge events time series may be driven by the subset of edges with a more canonical response function. Future work can compare different response functions, including agnostic finite impulse regressors, to see which produces the most reliable results.

Edge time series are also noisier than ROI time series. Therefore, future work should examine whether a minimal amount of smoothing might reduce noise and improve estimates of deflections. Applying a Gaussian smoothing kernel begins to approximate estimating dynamic functional connectivity using a sliding Gaussian window, but these approaches are not identical, as edges are computed using the mean and standard deviation of the ROIs from the whole time series, so information from outside of the Gaussian window is still being incorporated.

This work also compared max-T and NBS thresholding approaches for identifying significantly deflected edges. Max-T was quite conservative, identifying few edges consistent across datasets. In contrast, NBS identified a reliable set of edges for every contrast where a significant subcomponent was identified in each dataset. However, the overlap found between datasets was relatively small, suggesting that NBS may not be conservative enough. NBS is a cluster correction approach, which means that, like all cluster-based approaches, the significant subcomponents are often inflated by false positives. At best, a significant subcomponent can be interpreted as containing at least one significant edge (Sassenhagen & Draschkow, 2018, Rosenblatt et al., 2018). We chose a significance threshold of .01, in keeping with the creators of NBS (Zalesky et al., 2010, Serin et al., 2021), but it is worth noting that this may still produce many false positives. Other work examining cluster correction methods in activation analyses has suggested a threshold of .001 (Woo et al., 2014). Methods which reduce the possibility of false negatives *and* false positives when working with edges will be highly valuable.

We circumvented these issues by focusing on edges which were identified as members of the significant NBS components in both datasets. This required that a given edge survive thresholding (p < .01) and be adjacent to enough other significant edges, in both datasets, effectively producing a final threshold that is more stringent than p < .0001. Other research groups could replicate this approach either by using open data which matches the task they are studying, or through a split-half approach within their dataset (provided it is large enough).

Selecting edges based on an effective threshold more stringent than *p* < .0001 is likely unnecessarily conservative. An additional or alternative avenue consists of identifying alternatives to NBS that may better control for false positives. One option is the degree-based statistic (DBS; Yoo et al., 2017). This approach identifies clusters as edges sharing a single node, supporting some level of spatial inference, and examines them across a range of thresholds to assess significance. Another alternative is All-Resolutions Inference (ARI; Rosenblatt et al., 2018). ARI is a cluster correction approach that attempts to control the exact proportion of false positives within each cluster. While it has not been applied to connectomes to our knowledge, it relies on a notion of adjacency to identify clusters, and this could be translated into graphs (i.e., two edges are adjacent if they share a node). Future work can compare these methods to assess their sensitivity and ability to control for false positives.

We have shown that a simple modification to traditional fMRI analyses, converting ROI activity into edge cofluctuation time series, allowed us to extract novel and reliable information in the form of high-frequency edge deflections. One of the major benefits of this approach is its ability to produce windowless estimates of network strength in order to identify rapid network reconfigurations. In some ways, this is comparable to change point detection (e.g., Xu and Lunquist, 2015) and latent state analyses, which identify a set of brain states based on ROI activation and covariance that can be assigned to each time point in an fMRI time series. These analyses are in a sense windowless as they can identify shifts between brain states that happen from one moment to the next. Such analyses have identified large scale network changes tied to changes in attention (Yamashita et al., 2021; Cai et al., 2021; Kondo et al., 2022; Song et al., 2022). We believe this edge-based approach is complementary to latent state analyses in two ways. The first is that it can easily be combined with temporal regressors to extract information from edge cofluctuations. In contrast, latent state models are typically entirely data driven, and states must be linked to temporal events after the fact. The second is that the edge-based approach operates at a higher-dimensional description than whole-brain state changes, which describe coarse, low dimensional changes. For example, a study may involve participants switching between two attention demanding tasks that are highly similar and only differ in a small set of cognitive operations (for example, hearing versus reading some linguistic stimuli). It may be more conceptually useful to consider subsets of regions changing their connectivity profile to deal with this change in stimulus modality (a small change in a high dimensional space), rather than thinking of the brain as oscillating between two similar but distinct brain states (a large change in a low dimensional space).

This edge-based approach directly incorporates the perspective of network neuroscience that the brain may be understood in terms of the subsets of regions coordinating to perform certain cognitive processes, while enabling a finer temporal granularity than previous approaches. As a result, this method can be applied to any research which seeks to apply the network neuroscience perspective to relatively fast-changing cognitive phenomena. For example, how do networks reconfigure to form “process-specific alliances” when engaging different components of processes during learning (Cabeza & Moscovitch, 2013)? How do networks reconfigure in response to prediction errors during reinforcement learning paradigms? Or, how do networks change in response to increasing working memory loads, especially at supracacity set sizes? A broad swath of questions such as these are well posed for this edge-based approach. We hope that this work, along with the openly available code and data, will allow researchers to identify rapid network reconfigurations underlying a diverse set of cognitive processes.

This edge-based approach has offered an expanded perspective on rapid network reconfigurations during attentional fluctuations. While the default mode and attentional networks were implicated in lapses and successful attentional control as expected, so were subcortical and cerebellar regions, which linked to both associative and sensory-motor regions. In addition, there is now the possibility to investigate attentional fluctuations in terms of the relative integration or segregation of a given network (such as the medial frontal network) with respect to the rest of the brain. In conclusion, this expanded perspective can allow us to identify both new connections and new graphical motifs underlying differences in attention across time.

## Acknowledgements

This research was supported by National Science Foundation BCS-2043740 to M.D.R., National Science Foundation BCS-1558497 and National Institutes of Health MH 108591 to M.M.C., and a Neubauer Family Foundation Distinguished Scholar Doctoral Fellowship from the University of Chicago to H.M.J. We thank John Veillette for a discussion of all-resolutions inference in response to a presentation of this work. We thank Andrew Hannum for his suggestions on visualizing the edge-level heatmaps (Figure S2). We thank Chris Markiewicz for his suggestions on surface projecting the ROI results.

## Code & Data Availability

The code for this analysis is available at github.com/henrymj/edge_GLM. Dataset 2 can be accessed at https://nda.nih.gov/edit_collection.html?id=2402. For inquiries about Dataset 1, contact the authors of the original manuscript https://doi.org/10.1038/nn.4179.

## SUPPLEMENTAL FIGURES

**Figure S1.**
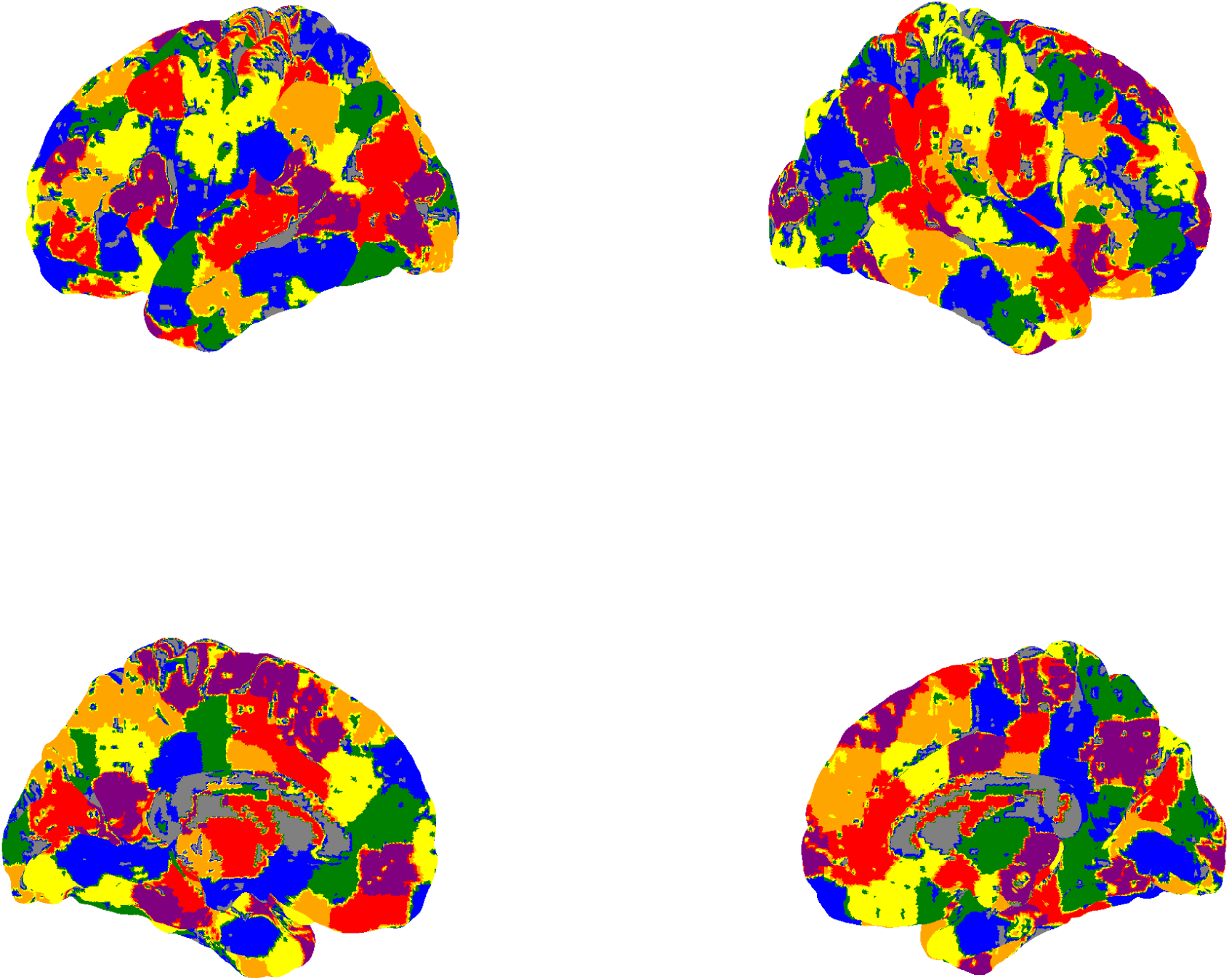
Shen atlas ROIs surface-projected with high contrast colors. Each ROI was mapped onto 1 of 6 highly contrasting colors around the color wheel (blue, green, yellow, orange, red purple) in a circularly repeating manner (e.g., both region 1 and region 7 were mapped onto blue, region 2 and region 8 were mapped onto green, etc). Each vertex was matched with the label of the nearest voxel.

**Figure S2.**
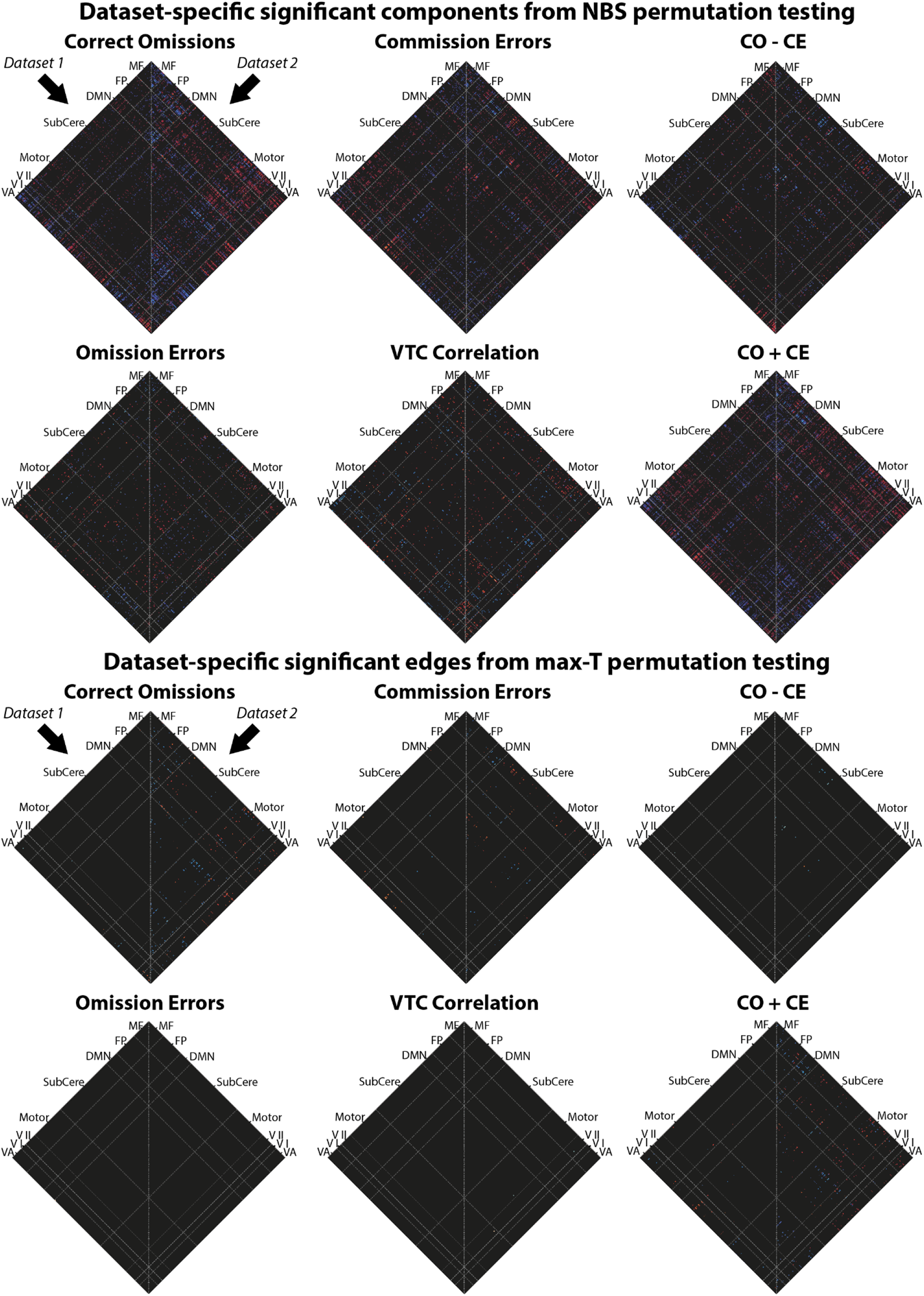
Dataset-specific second-level results for each of the five main contrasts and the correlation with the VTC. Within each results matrix, the left half reflects dataset 1 and the right half reflects dataset 2. Colors reflect thresholded t-scores, with blue indicating negative values and red indicating positive values. The top set reflects component thresholding using NBS permutation testing while the bottom reflects edge level thresholding using max-T permutation testing.

**Figure S3.**
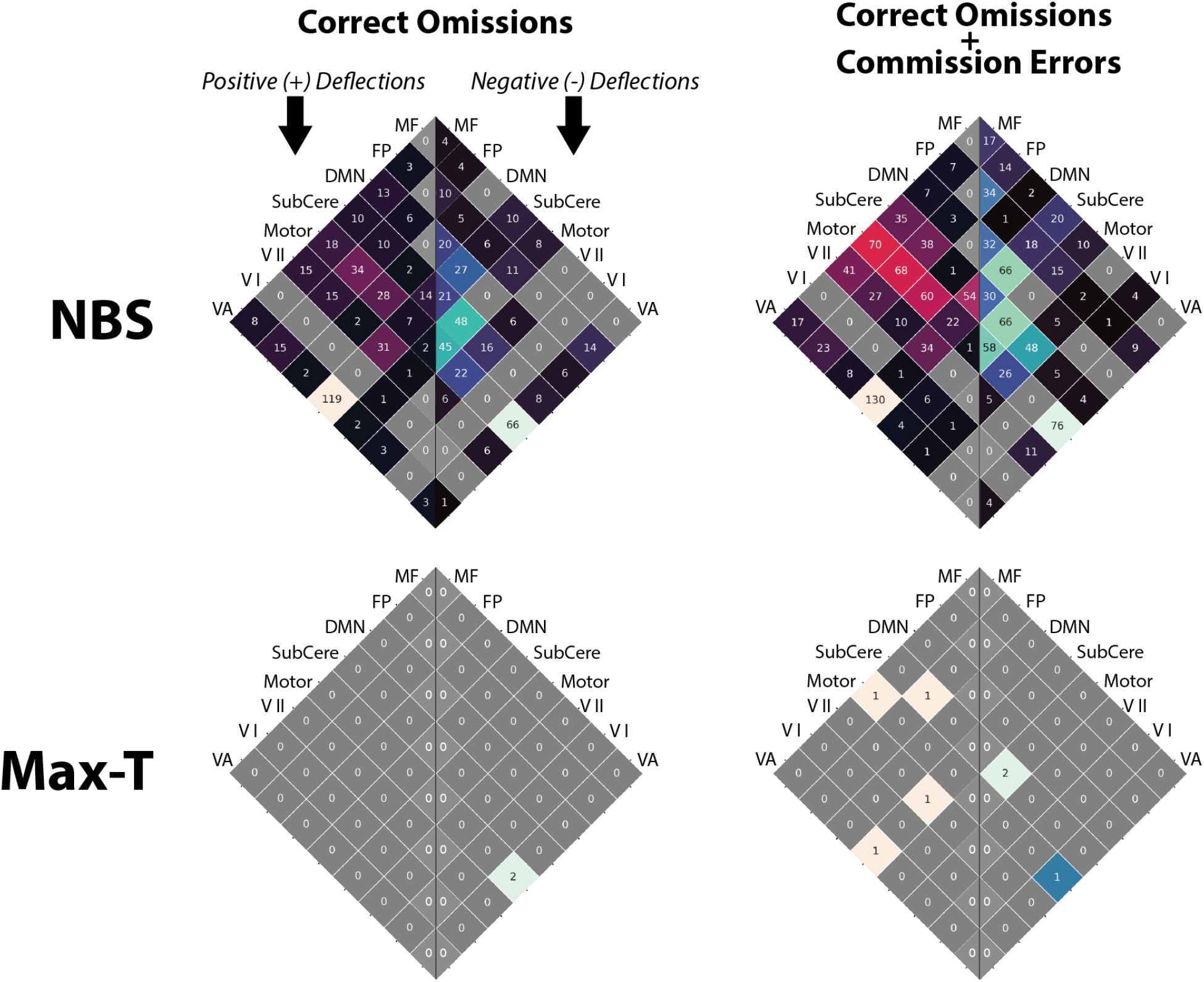
Cross-network distribution of reliable (i.e., significant in both datasets) edges from trial-type contrasts. Each heat map shows the number of edges in a given set of network-network connections that were significant for a given contrast (column). The top row reflects the overlap of components identified as significant via NBS permutation testing The bottom row reflects overlap of edges identified as significant via max-T permutation testing;. Note that for the Correct Omissions vs Baseline contrast (top left), the NBS overlap matrix is the same as that in Figure 2.

**Figure S4.**
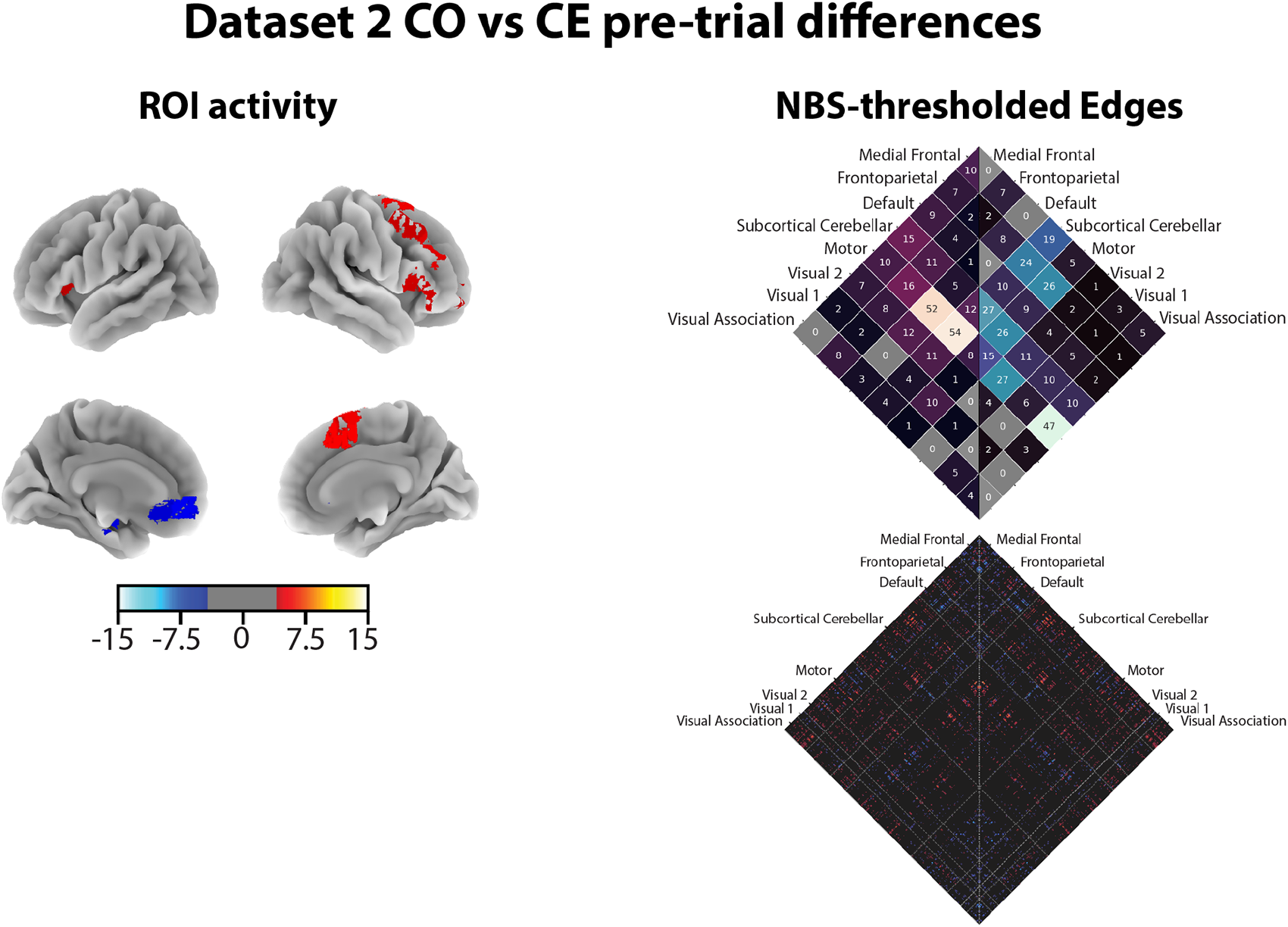
Summary of results for CO-CE trial precursor analysis in dataset 2. The left column shows the surface projection of the ROI activation results. The right column, upper matrix shows the count of significant edges, thresholded using NBS-based permutation testing, for each set of network connections. The left half reflects significantly positive edge counts, while the right reflects significantly negative edge counts. The lower matrix shows each significant edge’s t-score, with red reflecting positive values and blue reflecting negative values.

## Footnotes

1 Our data and preprocessing steps differ from that of Fortenbaugh et al. (2018) in a few ways. For example, Fortenbaugh et al. (2018) regressed out the mean signal from a white matter and CSF mask, while Rosenberg et al. (2016) and Yoo et al. (2022) regressed out the global signal, which includes grey matter. Therefore, this analysis also assesses whether the results of Fortenbaugh et al. (2018) are robust to processing differences.

2 We did not mention OE against baseline in our pre-registration, but we included the contrast to help ground our replication of Fortenbaugh et al. (2018).

3 In the preregistration, we also planned to fit a similar model, but with the VTC demeaned before convolution. However, this transformation only changes the beta values of the unmodulated event regressors, not that of the VTC regressor. Because we constrain our focus to the VTC results to compare with previous literature (Fortenbaugh et al., 2018, Esterman et al., 2013), we have omitted the demeaned approach.

4 Although we preregistered .05 as our initial threshold, .01 is more in keeping with other work using NBS (e.g., Zalesky et al., 2010, Serin et al., 2021), and should reduce the number of false positives in the final components, making them more interpretable.

5 Here we note two differences from our preregistration. First, the permutation testing approach differs from that described in our preregistration, which involved assessing the overlap of significant edges between datasets by assuming a hypergeometric distribution. The non-parametric permutation testing approach reduces the number of assumptions required when assessing significance with this new method. Second, in our preregistration we described assessing which networks contained more significant edges than expected by chance and comparing the overlap of networks across datasets. However, this approach is redundant with testing the overlap of edges across datasets and describing the distribution of overlapping edges across networks.

6 Whereas Esterman et al. (2013) observed more DMN and less dorsal attention network (DAN) activity before commission errors, Fortenbaugh et al. (2018) replicated the pre-trial DMN but not DAN effect. We did not formally test for replication of these network-level effects as our analyses were performed at the ROI level using the Shen 268-node atlas. Shen atlas nodes have been grouped into a DMN mask but not a DAN mask.

