## Supplemental Figures for "Edge-based general linear models capture high-frequency fluctuations in attention"

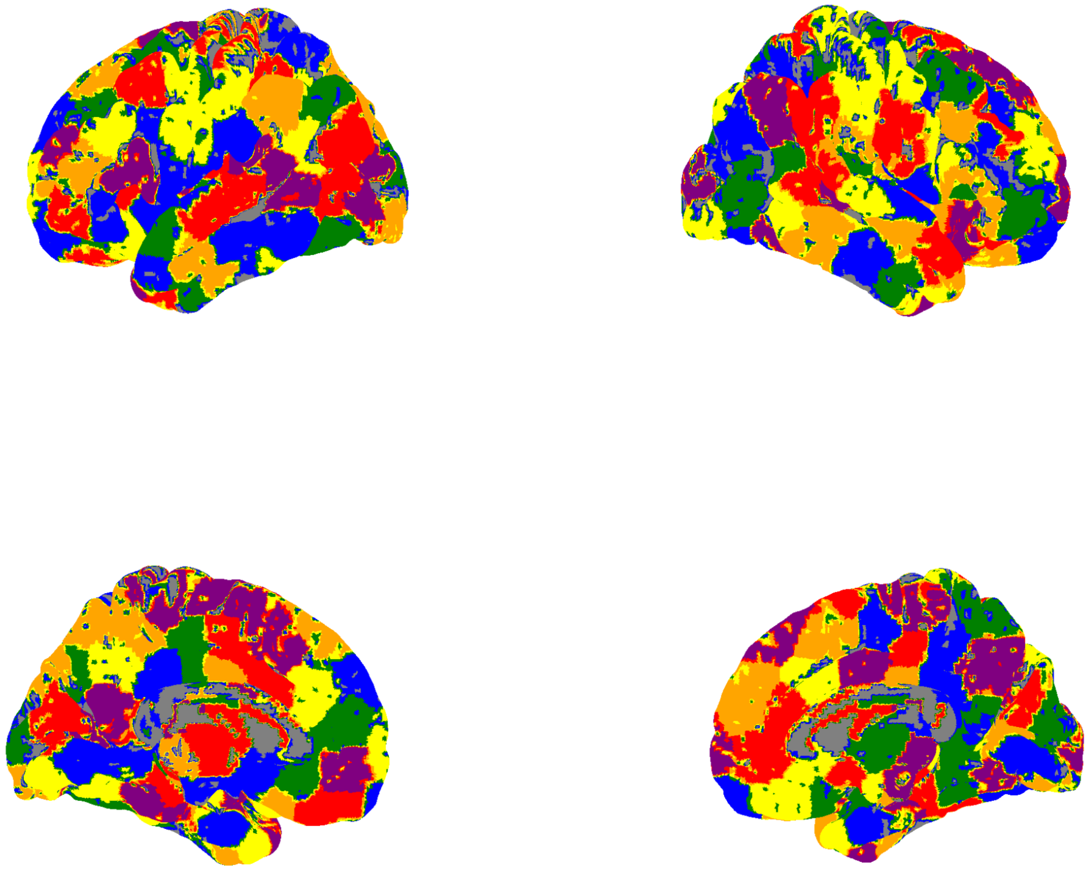

***Figure S1.*** *Shen atlas ROIs surface-projected with high contrast colors. Each ROI was mapped onto 1 of 6 highly contrasting colors around the color wheel (blue, green, yellow, orange, red purple) in a circularly repeating manner (e.g., both region 1 and region 7 were mapped onto blue, region 2 and region 8 were mapped onto green, etc). Each vertex was matched with the label of the nearest voxel.*

### Dataset-specific significant components from NBS permutation testing

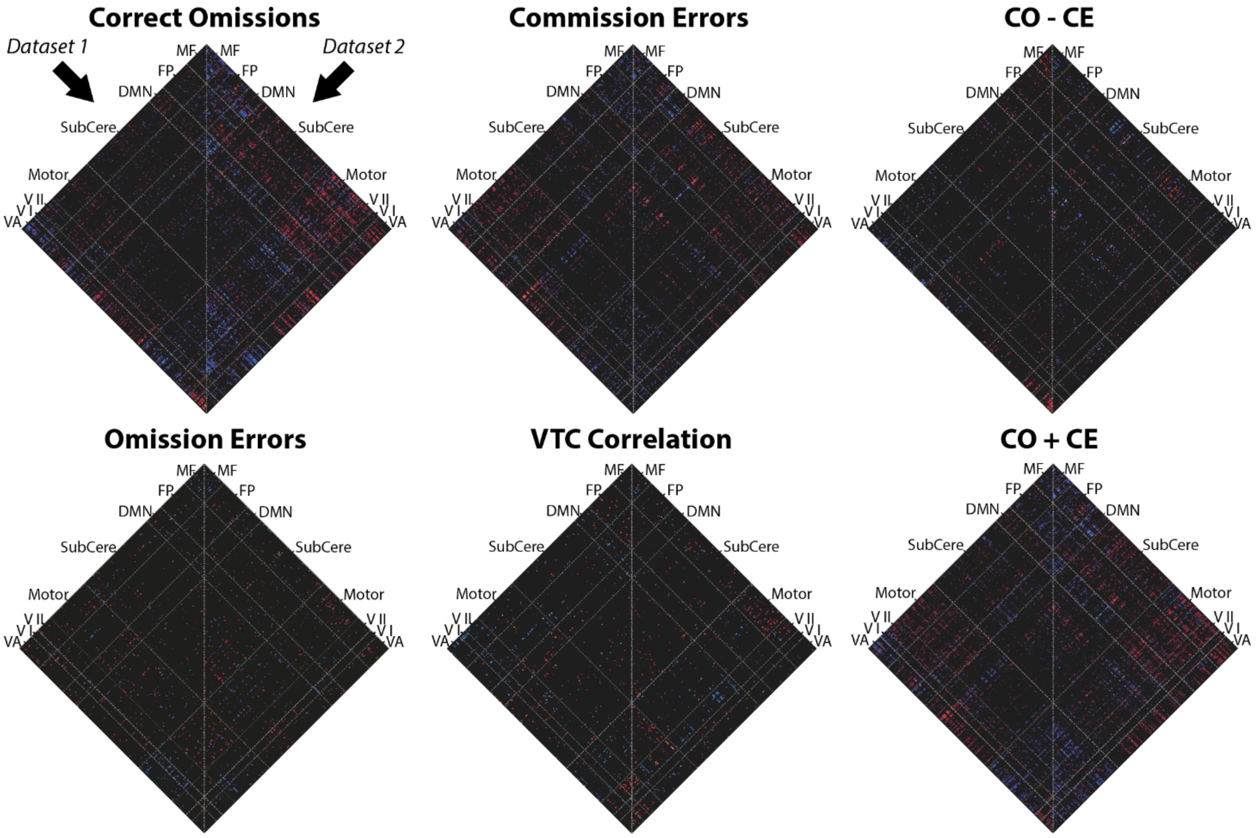


### Dataset-specific significant edges from max-T permutation testing

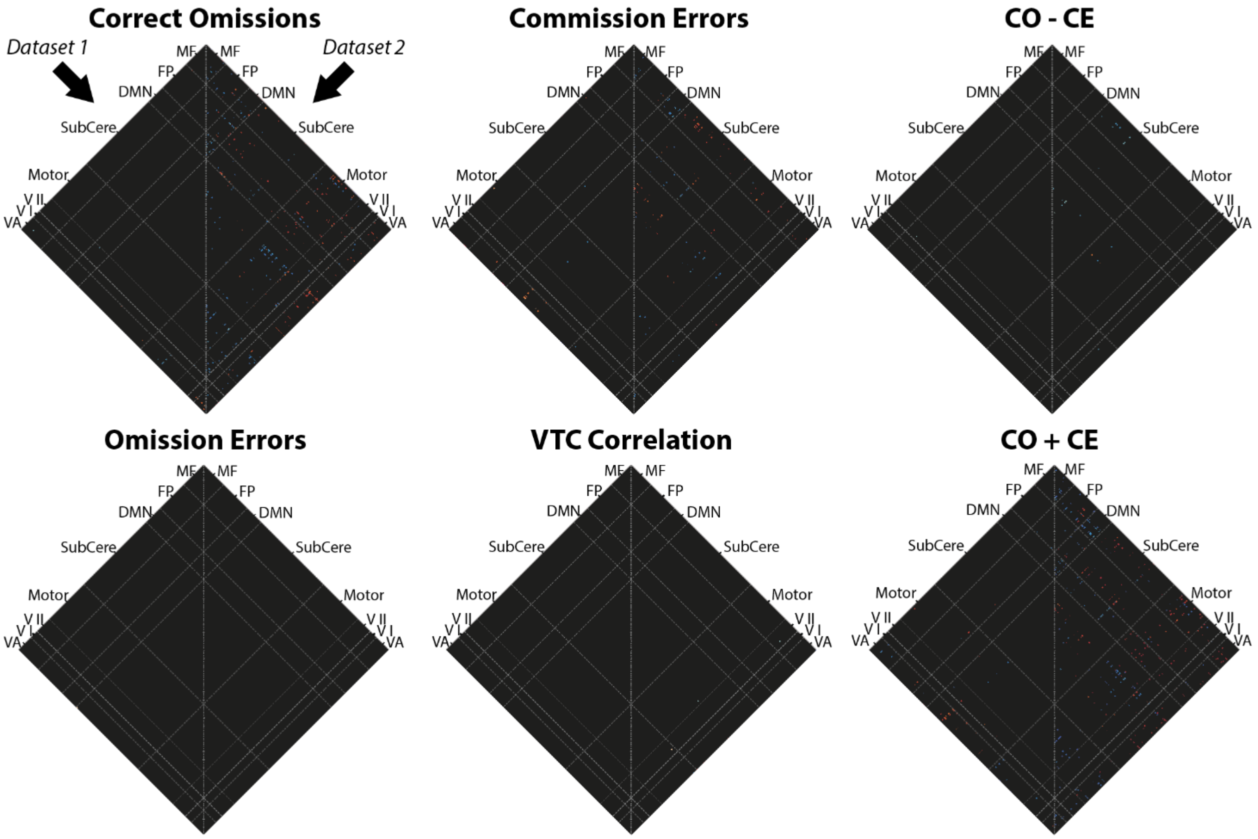

***Figure S2.*** *Dataset-specific second-level results for each of the five main contrasts and the correlation with the VTC. Within each results matrix, the left half reflects dataset 1 and the right half reflects dataset 2. Colors reflect thresholded t-scores, with blue indicating negative values and red indicating positive values. The top set reflects component thresholding using NBS permutation testing while the bottom reflects edge level thresholding using max-T permutation testing.*

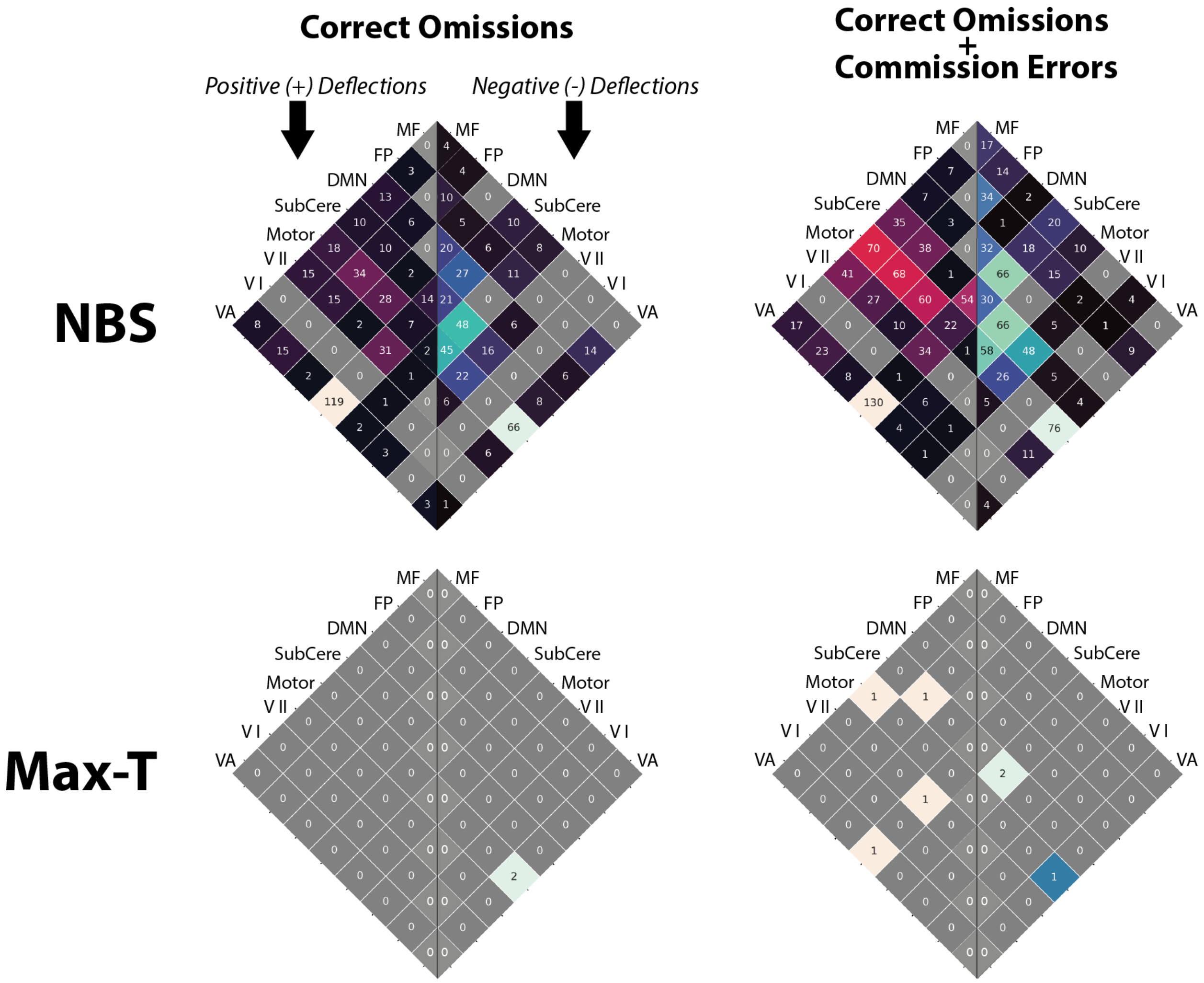


***Figure S3.*** *Cross-network distribution of reliable (i.e., significant in both datasets) edges from trial-type contrasts. Each heat map shows the number of edges in a given set of network-network connections that were significant for a given contrast (column). The top row reflects the overlap of components identified as significant via NBS permutation testing The bottom row reflects overlap of edges identified as significant via max-T permutation testing;. Note that for the Correct Omissions vs Baseline contrast (top left), the NBS overlap matrix is the same as that in Figure 2.*

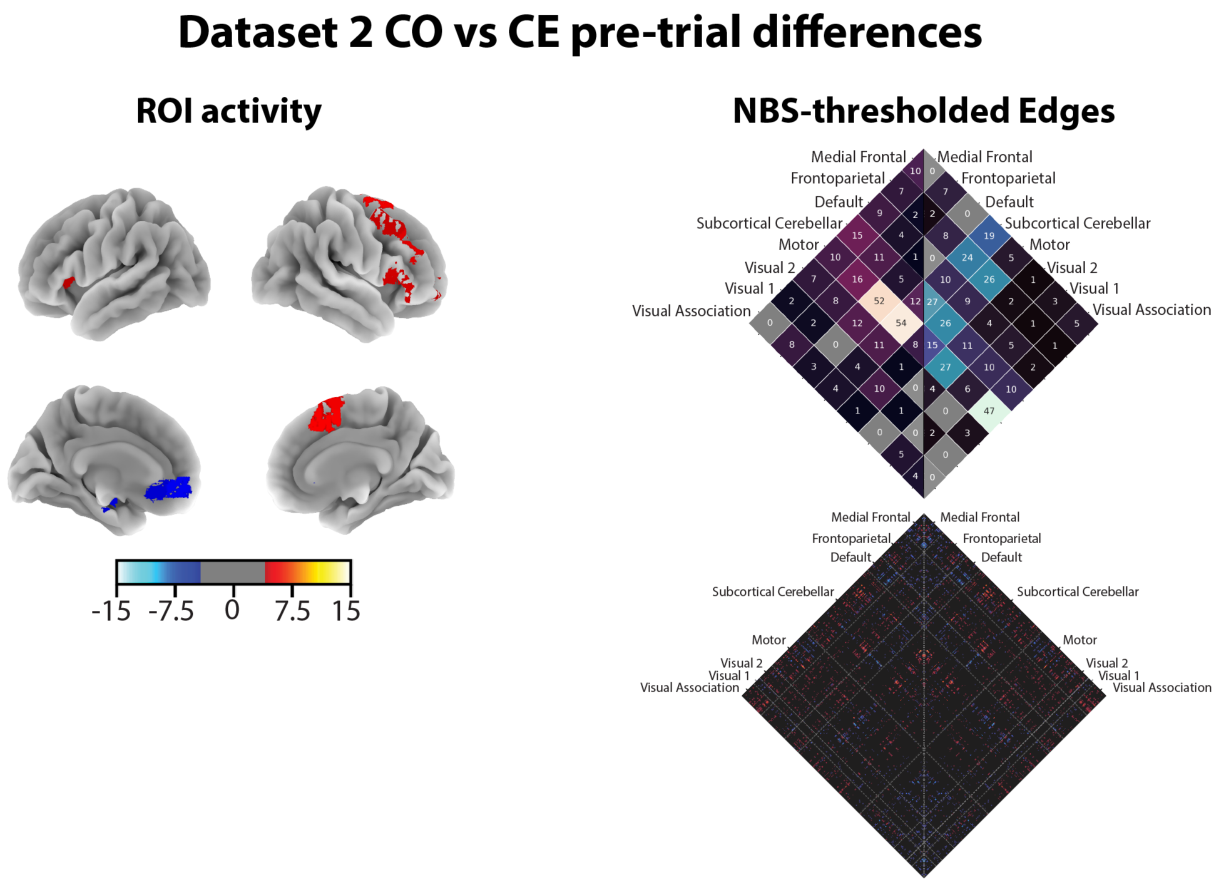


***Figure S4***. Summary of results for CO-CE trial precursor analysis in dataset 2. The left column shows the surface projection of the ROI activation results. The right column, upper matrix shows the count of significant edges, thresholded using NBS-based permutation testing, for each set of network connections. The left half reflects significantly positive edge counts, while the right reflects significantly negative edge counts. The lower matrix shows each significant edge's *t*-score, with red reflecting positive values and blue reflecting negative values.
